# Macrophage signaling history constrains subsequent transcriptional responses to cytokine stimulation

**DOI:** 10.64898/2026.11.14.765729

**Authors:** Jacob B. Hochfelder, Rachel A. McGinn, Cal Gunnarsson, Diana N. Gong, Bryan D. Bryson

**Affiliations:** Department of Biological Engineering, Massachusetts Institute of Technology, Cambridge, MA, USA; Ragon Institute of MGH, MIT, and Harvard, Cambridge, MA, USA

## Abstract

Macrophages perform a wide array of distinct and crucial functions, and therefore tightly regulate their responses to external stimuli. These cells must integrate sequential signals over time to mount an appropriate response to their immunological niche. How prior cytokine stimulation shares macrophage responses to a subsequent stimulus remains poorly understood, yet has important implications for pathogen clearance as well as tissue repair and homeostasis. Here, we systematically characterize transcriptional responses to sequential cytokine stimulation in primary human macrophages, as well as demonstrate the effect of interactions with bacterial pathogens on macrophage transcriptional responses and plasticity. We find that prior stimulation history constrains subsequent transcriptional responses in a stimulus-specific and asymmetric manner. Across the cytokines we tested, we observe that IFNβ is a dominant regulator of subsequent responses to other cytokines. We observe that this plasticity restriction is concentration dependent, rapidly established, and persists after cytokine removal. Additionally, we show that endogenous IFNβ produced during infection with *Mycobacterium tuberculosis* restricts subsequent cytokine responses in a manner that is dependent on the activity of the bacterial secretion system ESX-1. Taken together, these findings establish a hierarchical model of macrophage responses to cytokines that depends on cytokine identity and order.

## Introduction

Macrophages are key players of the immune system during both homeostasis and disease, and must integrate many signals over time to coordinate their responses^1^. These cells exhibit notable functional diversity, from phagocytosis of pathogens to tissue remodeling^2,3^. As a result, macrophages synthesize information from extracellular signals to adopt distinct transcriptional and functional activation states^1^. The model of activation states has expanded beyond the traditional M1/M2 classification^4^ to include a more complex parameter space of functions and responses^5–8^. There is a long-standing paradigm that monocyte-derived macrophages are highly plastic, meaning that they can adopt multiple activation states over their cellular lifetimes^3,9^. This plasticity is necessary for resolving inflammation after infection, adapting to new pathogen challenges, and remodeling the tissue microenvironment^10^.

However, recent studies have challenged this model, instead demonstrating that prior activation can constrain the transcriptional response to subsequent stimuli^11–13^. These plasticity restrictions can have durable effects on immune responses due to the longevity of recruited macrophages in tissues after monocyte differentiation^10,14^.

Previous work has described limitations to macrophage plasticity, both at the level of global cellular state and through specific signaling mechanisms. One commonly used experimental design involves pretreating macrophages with interferon-γ (IFNγ), a Th1 cytokine, and lipopolysaccharide (LPS), a Toll-like receptor (TLR) ligand, to trigger a traditional inflammatory state, then restimulating the cells with interleukin-4 (IL-4) to understand how IL-4 signaling is affected by pretreatment compared to the naive response. In bone-marrow derived mouse macrophages, upregulated activity of inducible nitric oxide synthase (iNOS) after IFNγ and LPS stimulation disrupts oxidative phosphorylation normally associated with IL-4 signaling, preventing macrophages from adopting an IL-4 activation state^11^. Additionally, the durability of transcriptional and epigenetic cytokine responses depends on the stage of the cell cycle at the time of stimulation for mouse macrophages^15^. While these specific mechanisms of plasticity restrictions have been proposed in mouse macrophages, they are incomplete for human macrophages, which do not express iNOS^16^ or proliferate *in vitro*. Human macrophages have been shown to retain their stimulation history through chromatin remodeling, which both potentiates and diminishes subsets transcriptional responses to subsequent TLR ligand stimulation^12^. Furthermore, cytokines have recently been shown to remain on the cell surface resulting in prolonged signaling after wash-out^17^ (which has been observed in other contexts^18^). In addition to these global changes to cellular state, overlapping use in signaling machinery between two cytokines can result in negative feedback regulatory mechanisms that disrupt responses to subsequent signaling events^19^. Additionally, stimulating macrophages with opposing signals can result in heterogeneous effects within a single population of cells^20^, or dynamic transitions in macrophage state due to differences in signaling kinetics between cytokines^21^. This diversity in plasticity restriction mechanisms highlights the complexity through which macrophages integrate multiple conflicting signals over time. However, prior work has largely focused on individual cytokine pairs, leaving unaddressed how plasticity restrictions operate across a broader landscape of activation states in primary human macrophages.

Here, we repolarize macrophages by stimulating them with one cytokine, then restimulating them with a different cytokine. We model a diverse set of signaling sequences by using recombinant cytokines to allow for experimental control of the identity, order, concentration, and duration of stimuli, each of which has well-characterized responses^6,22^.

Using a multi-modal approach, we employ transcriptomics, protein measurements, epigenomics, and synthetic chimeric receptors to demonstrate sequences of cytokines that exhibit plasticity restrictions. We find that IFNβ dominates macrophage transcriptional state across many other cytokines in an asymmetric and tunable manner that is mediated through IFNAR2/JAK1 signaling. We further characterize the dynamics and concentration dependence of type I interferon plasticity restrictions. Finally, we show that endogenous IFNβ production, resulting from the activity of the *Mycobacterium tuberculosis* (Mtb) secretion system ESX-1, restricts subsequent IL-4-induced gene expression, demonstrating that plasticity restrictions operate in a physiologically relevant infection context.

## Results

### Macrophage transcriptional state reflects both prior and current cytokine stimulation

To understand how sequential cytokine stimulation informs macrophage state, we cultured primary human monocyte-derived macrophages (hMDMs) with all pairwise permutations from a panel of six cytokines. Cytokines was chosen to represent a broad range of macrophage states and functions, as well as relevance in the context of infection: IFNβ, IFNγ, IL-1β, IL-4, IL-10, and transforming growth factor beta (TGFβ) (Figure 1B). IFNβ and IFNγ are potent drivers of inflammatory immune responses to viral and bacterial infections, primarily through STAT1-mediated transcriptional responses^23^. Macrophages both produce and respond to IL-1β, resulting in NF*κ*B signaling and increased inflammatory chemokine production^24^. IL-4- and IL-10-stimulated macrophages upregulate gene expression programs related to tissue repair and resolution of inflammation primarily through STAT6 and STAT3 phosphorylation, respectively^25,26^. TGFβ signaling induces SMAD phosphorylation and upregulation of extracellular matrix remodeling programs in macrophages^27^. Macrophages were cultured with each cytokine at 10 ng/mL, which robustly activated the expected downstream signaling pathways (Supplementary Figure 1). We included all forward and reverse pairwise cytokine combinations to understand the effect of each combination and stimulation order (Figure 1B). Macrophages were cultured with the first cytokine for 24 hours, washed three times with PBS, and then stimulated them with a second cytokine for an additional 24 hours before performing bulk RNA sequencing (Figure 1C). In addition to unstimulated controls, we included conditions in which macrophages were stimulated with a single cytokine during only the first 24 hours, only the second 24 hours, or continuously for 48 hours to account for transcriptional differences associated with stimulation duration and timing. PBMC donors were distributed across culture plates and library preparation batches to minimize batch effects.

**Figure 1:**
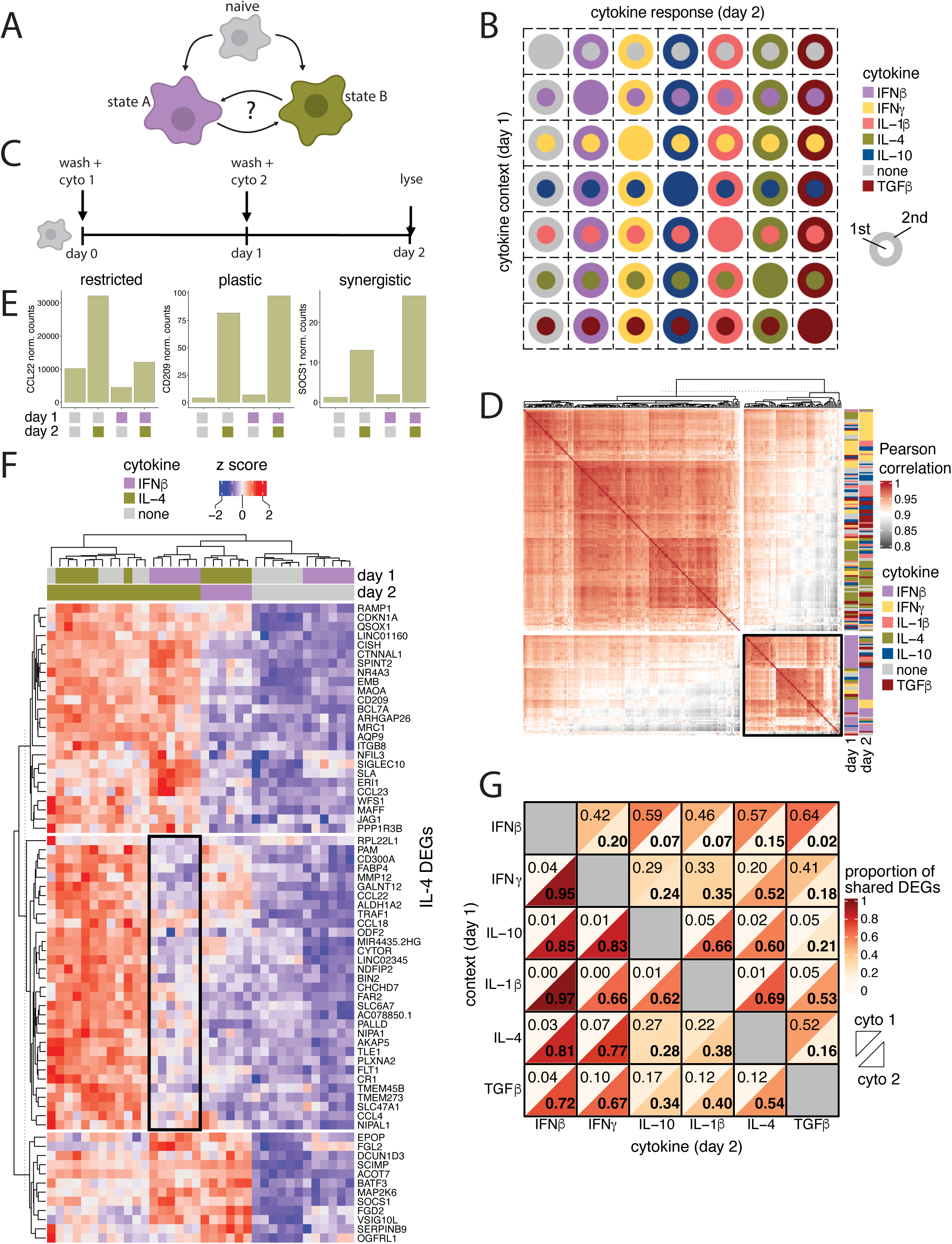
Sequential cytokine stimulation reveals plasticity restrictions in macrophage transcriptional state. **A)** Model of macrophage repolarization as subsequent stimulation after initial polarization. **B)** Schematic of stimulations performed. **C)** Experimental set-up. N = 6 PBMC donors x 2 technical replicates of library prep performed on different days. **D)** Pearson correlation of normalized counts matrix after RNA sequencing. K means clustering (k=2) shows IFNβ-dominated cluster, highlighted with black box. **E)** Barplots of normalized counts of IL-4 differentially expressed genes that are restricted, plastic, or synergistic effects from IFNβ pretreatment. **F)** Heatmap of IL-4 differentially expressed genes, with conditions containing only IFNβ, IL-4, or none shown, k means clustered (k=3). Subset of IL-4 DEGs that are sensitive to IFNβ prestimulation, highlighted with black box. **G)** Heatmap of proportion of DEGs relative to unstimulated condition. Upper left triangle represents the proportion of genes that are DE in repolarized condition that are also DE with the first cytokine alone. Lower right triangle represents the proportion of genes that are DE in repolarized condition that are also DE with the second cytokine alone.

After stimulations, each sample was lysed and prepared for bulk RNA sequencing to characterize the effect of each cytokine sequence on overall transcriptional state. First, we calculated the Pearson correlation across all samples to understand if sequentially stimulated macrophages adopted transcriptional states more similar to the first or second cytokine. Under a model of unrestricted plasticity, macrophages stimulated with the same second cytokine would be expected to share similar transcriptional states regardless of prior stimulation history. Instead, transcriptional similarity between conditions was better predicted by the identity of the first cytokine than the second for all IFNβ-pretreated conditions and a subset of IFNγ and IL-4 pretreated conditions (Figure 1D). This suggests that prior cytokine stimulation resulted in restricted plasticity in the subsequent macrophage transcriptional response. These effects were asymmetric across cytokine pairs. In particular, IFNβ almost exclusively formed a unique cluster irrespective of cytokine order. To examine these effects at the gene level, we focused on IFNβ and IL-4 as a model cytokine pair. These two cytokines drive highly distinct transcriptional programs. IFNβ drives STAT1-mediated inflammatory gene expression^23^ and IL-4 drives

STAT6-mediated tissue repair programs^25^. Additionally, both cytokines produced the largest numbers of DEGs in our dataset, making their interaction an informative model for understanding how prior stimulation constrains the transition to a distinct second activation state. To answer this question, we analyzed IL-4-induced DEGs when macrophages were treated with IL-4 with or without IFNβ. We observed genes that were upregulated by IL-4 alone but no longer upregulated with IFNβ pretreatment (ex. CCL22), genes that were unaffected by IFNβ pretreatment (ex. CD209), and genes that exhibited increased expression with both IFNβ and IL-4 (ex. SOCS1) (Figure 1E). Almost half (46%) of these DEGs formed a plasticity restricted cluster: naive macrophages upregulate them in response to IL-4, but failed to do so following IFNβ pretreatment (Figure 1F). This plasticity-restricted subset included chemokines (*CCL22*, *CCL18*, *CCL4*), as well as genes related to lipid metabolism and energy production (*FAR2*, *FABP4*, *CHCHD2*), and cytoskeletal organization (*CYTOR*, *PALLD*, *BIN2*). The remaining genes were induced by IL-4 independent of IFNβ pretreatment, or their expression was higher in conditions with both IFNβ and IL-4 than IL-4 alone.

Having established that IFNβ pretreatment restricts a subset of IL-4-induced genes, we next asked more broadly whether the transcriptional state of repolarized macrophages more closely resembles the first or second cytokine. To answer this, we asked what fraction of DEGs in each repolarized condition were also differentially expressed in response to either cytokine alone. For each repolarized condition, we compared its DEGs against those induced by each single cytokine stimulation. For example, for the IFNβ → IL-4 condition, we asked what proportion of DEGs were also seen with IFNβ alone versus IL-4 alone (Supplementary Figure 2). We excluded genes that were differentially expressed in response to both single cytokines, since these could not distinguish the influence of one cytokine from the other. In repolarized conditions where IFNβ was in the first cytokine, the majority of DEGs were shared with IFNβ alone rather than the second cytokine (Figure 1G), indicating that IFNβ pretreatment dominates the overall transcriptional state even after an intervening stimulus. In contrast, repolarized conditions where IFNγ was the first cytokine showed a more balanced mixture of IFNγ-associated DEGs and DEGs associated with each second cytokine, suggesting that IFNγ-primed macrophages more readily integrate a subsequent signal. Together, these data reveal a difference between type I and type II interferons in how durably they shape macrophage transcriptional state. IFNβ maintains a dominant transcriptional imprint over an intervening stimulation event, while IFNγ permits more balanced co-expression of programs from both cytokines.

### Preexisting macrophage transcriptomic state influences ability to repolarize

Differential expression analysis of repolarized macrophages revealed sets of genes that are confidently upregulated for each cytokine, but strict p-value and log fold change cutoffs may have obscured subtle changes in gene expression. To assess global transcriptional similarity across conditions without the constraints of significance cutoffs, we performed principal component analysis (PCA). Strikingly, conditions stimulated with IFNβ clustered tightly along PC1 and conditions stimulated with IL-4 clustered tightly along PC2 (Figure 2A). Genes associated with IFNβ responses contributed strongly to PC1 (*IFI27*, *IFITM3*, *RSAD2*), while genes associated with IL-4 responses contributed strongly to PC2 (*SCIMP, FABP4*, *CCL22*). (Supplementary Figure 3B). Additionally, we observed that many conditions clustered by the cytokine in the first 24 hours when that cytokine was IFNβ or IL-4, suggesting that the macrophage state is still influenced by these cytokines after an intervening signal.

**Figure 2:**
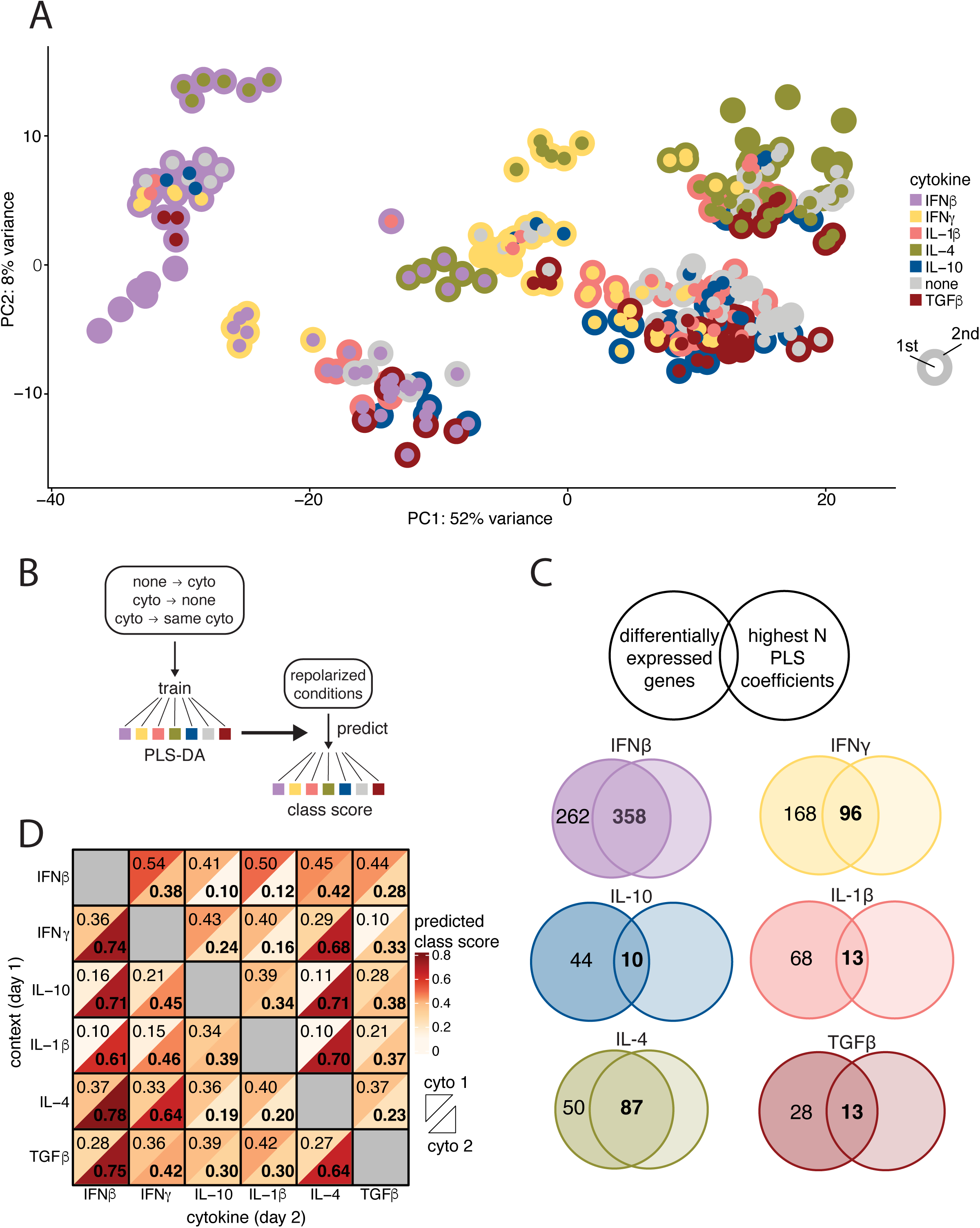
Cytokine regulation of gene expression programs is influenced by a mixture of current and previous stimuli **A)** Principal component analysis on dataset, PC1 and PC2 shown. Each dot is one biological replicate, technical replicates combined. **B)** Schematic of PLS-DA model. Model was trained on all donors and all conditions with a single cytokine identity (day 1 only, day 2 only, and day 1 plus day 2). **C)** Comparison of genes from DE analysis versus PLS-DA. Left circles represent genes that are differentially expressed relative to unstimulated. Right circles represent the top “n” genes by their coefficient for each PLS class, where “n” is determined by the number of differentially expressed genes for that cytokine. **D)** Predicted class scores from PLS-DA model for each repolarized condition split by each cytokine present on that condition. Upper left triangle represents cytokine 1, lower right triangle represents cytokine 2.

Beyond assessing transcriptional similarity across conditions with unsupervised methods, we additionally tested whether cytokine stimulation history could be inferred from macrophage transcriptional state. To do this, we used partial least squares discriminant analysis (PLS-DA) to distinguish transcriptional responses by treating each cytokine as a class in the model. The PLS-DA model was trained on all conditions with a single cytokine (“none → cytokine X”, “cytokine X → none”, “cytokine X → cytokine X”), then applied to all repolarized conditions where two unique cytokines were present (Figure 2B). We observed some agreement between DEGs for each single cytokine stimulation and the genes with the highest feature weights in the PLS-DA model, but there were many distinct genes for every cytokine (Figure 2C). This is likely due to the fact that the PLS-DA prioritizes genes that discriminate classes in order to assign prediction confidence. The model was then applied to each repolarized condition and used to predict the scores for each of the 6 cytokines in the panel, in addition to a no stimulation score. For every sample where IFNβ was the first cytokine, the model assigned a higher score to the IFNβ class than the other cytokine in that condition despite the fact that other cytokine was the more recent stimulus (Figure 2D), suggesting that IFNβ dominates macrophage state even when it is no longer present. In contrast, many other repolarized conditions received similar scores for both cytokines, suggesting that both stimuli contributed substantially to the resulting transcriptional state. Overall, the PLS-DA generally assigned higher scores to cytokines that were present in each condition, demonstrating the specificity of each class in the model (Supplementary Figure 4). Taken together, these results suggest that macrophages integrate sequential stimulation information by expressing a combination of gene expression programs, but the relative contribution of each cytokine on total macrophage state depends on the identities of each cytokine.

### Interferon induced plasticity restriction occurs quickly and at relatively low concentrations

Previous work has shown that IL-4-induced surface marker expression can be disrupted by prior inflammatory stimulation^11^, leveraging flow cytometry as a relatively fast readout for assessing plasticity restrictions at the protein level. We therefore asked whether IFNβ pretreatment similarly restricts IL-4-induced surface marker expression in primary human macrophages, using CD206 as a readout due to its robust and consistent induction by IL-4 in hMDMs compared to other markers we tested (Figure 3B). Having confirmed that IFNβ → IL-4 treatment restricts CD206 upregulation, we then used this readout to systematically characterize how the concentration and duration of IFNβ pretreatment influence the extent of the plasticity restriction with respect to IL-4. We hypothesized that both variables would influence the extent of restricted plasticity, so we treated hMDMs with IFNβ at 10 ng/mL, 100 pg/mL, and 1 pg/mL for 1, 6, or 24 hours before washing and restimulating with IL-4 (Figure 3A). We observed the same diminished CD206 response at 1 hour of IFNβ treatment as 24 hours for macrophages pretreated with 10 ng/mL of IFNβ, suggesting that IFNβ rapidly establishes an impaired response to subsequent IL-4-induced CD206 expression (Figure 3B).

**Figure 3:**
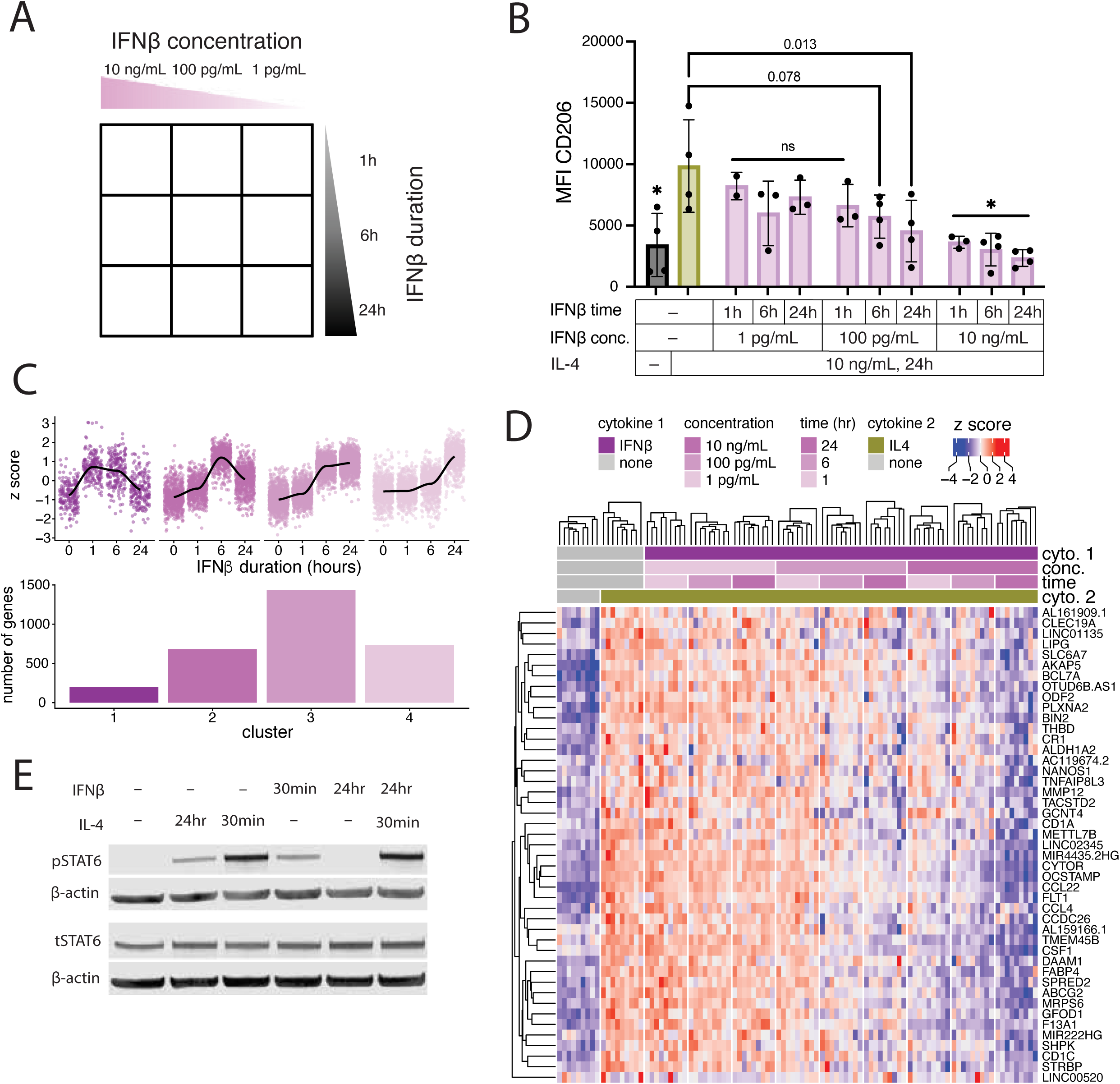
IFNβ-induced plasticity restriction is dependent on both concentration and duration of stimulation **A)** Schematic of conditions tested **B)** CD206 flow MFI reported above isotype control. N = 2-4 PBMC donors performed on 2-4 separate days. One-way ANOVA performed comparing all samples to IL-4, with Dunnett multiple hypothesis correction. Asterisks indicate p < 0.01. **C)** K means clustering (k=4) of 10 ng/mL IFNβ DEGs reveals transcriptional activity peaks before 24 hours. **D)** Normalized expression of IL-4 context dependent DEGs across IFNβ concentrations and times. Context dependent DEGs were selected based on IL-4 DEGs that are differentially expressed in IFNβ (24h, 10 ng/mL) → IL-4 relative to IL-4 alone. **E)** IFNβ does not affect IL-4-induced phosphorylation of STAT6. For **C** and **D**, RNA sequencing was performed on 6 PBMC donors with 2 technical replicates of library prep each, performed on 2 different days.

We also observed a similar plasticity restriction at 100 pg/mL of IFNβ at 24 hours. Shorter pretreatment durations with 100 pg/mL IFNβ progressively restored CD206 response to IL-4, suggesting that plasticity restrictions are tunable (Figure 3B). 1 pg/mL of IFNβ was not sufficient to establish the CD206 plasticity restriction at any timepoint. We note that *MRC1*, gene that encodes CD206, did not show restricted transcription following IFNβ pretreatment (Figure 1E) despite the reduction in CD206 surface protein as well as total cellular CD206 protein levels (Supplementary Figure 5A), suggesting the discrepancy occurs at a post-transcriptional level. While the mechanism underlying this transcript-protein discrepancy remains to be determined, CD206 surface expression has been previously shown as a reliable marker of plasticity restrictions in human macrophages^11^, and our transcriptomic data identify a broader set of plasticity-restricted genes beyond *MRC1* that more fully capture the scope of IFNβ-induced restriction.

Having established that IFNβ-induced plasticity restriction of CD206 expression is dependent on both concentration and duration of pretreatment, we next asked whether these same dependencies hold at the transcriptional level. To test this, we performed RNA sequencing on macrophages pretreated with IFNβ at each concentration and duration, with or without subsequent IL-4 stimulation. Before examining the IL-4 response, we first asked when IFNβ establishes transcriptional programs to understand whether the rapid plasticity restriction we observed at one hour could be explained by early IFNβ-induced gene expression.

Clustering all IFNβ-induced DEGs by their temporal expression pattern revealed that the majority of genes (2115/3051) were induced by 6 hours, either transiently or continuously through 24 hours (Figure 3C), indicating that the plasticity restriction is established before the peak of IFNβ transcriptional activity. The remaining genes were either transiently induced starting at 1 hour, or were not highly induced until 24 hours. We then examined whether the subsequent response of IL-4 induced DEGs showed the same concentration and duration dependence as CD206. Focusing on genes that were induced by IL-4 in naive macrophages but were plasticity restricted in macrophages pretreated with IFNβ at 10 ng/mL for 24 hours, we found that this transcriptional plasticity restriction was progressively less severe with lower IFNβ concentrations and shorter pretreatment durations (Figure 3D), mirroring our protein-level observations and confirming that IFNβ-induced plasticity restriction is tunable rather than a binary switch.

Due to the relatively fast establishment of plasticity restriction by IFNβ, we hypothesized that the initial IL-4 signaling cascade was affected by IFNβ pretreatment. To test this, we pretreated macrophages with IFNβ and then stimulated them with IL-4 for 30 minutes and looked for STAT6 phosphorylation via western blot since STAT6 is robustly phosphorylated upon IL-4 stimulation. We observed that 10 ng/mL 24 hour IFNβ pretreatment did not affect STAT6 phosphorylation, suggesting that the plasticity restrictions we observe are regulated downstream of early phosphorylation events after IL-4 stimulation (Figure 3E). We additionally tested whether this IFNβ-induced plasticity restriction is durable over time. Macrophages were stimulated with IFNβ for 24 hours, then washed and replaced with new media every day for an additional 72 hours before subsequent IL-4 stimulation. We observed the same plasticity restriction in CD206 expression with this 72 hour rest as we did without the rest, suggesting that this plasticity restricted state is long-lived (Supplementary Figure 5B). To test the hypothesis that epigenetic regulation plays a role in maintaining cell state and restricting transcriptional plasticity, we performed single-cell ATACseq on macrophages pretreated with IFNβ or IFNγ, then repolarized with IL-4. We cultured macrophages with 10 ng/mL of each cytokine for 24 hours and included DPBS washes as described above. Single stimulation and unstimulated conditions were also included as controls. We observed that IFNγ → IL-4 repolarized macrophages clustered with IFNβ and IFNγ single stimulation conditions while IFNβ → IL-4 repolarized macrophages clustered with either IL-4 or unstimulated conditions (Supplementary Figure 6A). This notably conflicted with our transcriptional observations, where we observe that IFNβ maintains a stronger plasticity restriction than IFNγ, suggesting that chromatin appears accessible for IL-4-induced programs that are transcriptionally restricted.

We additionally performed transcription factor (TF) motif enrichment analysis to understand which specific gene expression programs were affected by interferon pretreatment. We observed that IFNγ → IL-4 repolarized macrophages were highly correlated in TF motif enrichment with IFNγ-stimulated macrophages, suggesting that IFNγ pretreatment leaves a durable epigenetic imprint that persists after IL-4 restimulation (Supplementary Figure 6B). In contrast, IFNβ → IL-4 repolarized macrophages were highly correlated in TF motif enrichment with IL-4-stimulated macrophages and anticorrelated with IFNβ-stimulated macrophages.

Specifically, we observe maintained enrichment for proinflammatory TF motifs in the IFNγ → IL4 repolarized condition that were increased by stimulation with IFNγ alone, such as IRF9/8/2/7/1 and STAT1::STAT2 (Supplementary Figure 6C). Strikingly, we observe that these motifs are unenriched in IFNβ → IL-4 repolarized macrophages despite their enrichment in macrophages stimulated with IFNβ alone. This suggests that the effect of IL-4 stimulation on chromosomal organization is context dependent and influenced differently by type I and type II interferons. Additionally, this implies that IFNβ-induced plasticity restriction operates at a level beyond chromatin accessibility.

### IFNAR2/JAK1 signaling is sufficient to establish IFN№ plasticity restriction

To test potential mechanisms of IFNβ plasticity restrictions, we tested whether components of the IFNβ receptor heterodimer, IFNAR1 and IFNAR2, were enough to cause a plasticity restriction. Macrophages were pretreated with either deucravacitinib or upadacitinib before IFNβ stimulation. Deucravacitinib targets TYK2, which binds to the IFNAR1 subunit, and upadacitinib targets JAK1, which binds to the IFNAR2 subunit. To determine the efficacy of drug treatment in blocking IFNβ treatment in vitro, we measured pSTAT1, which becomes highly phosphorylated after IFNβ stimulation. Macrophages were treated with 100 times each molecule’s respective IC50 (4.3 µM for upadacitinib and 100 nM for deucravacitinib) for one hour minutes before stimulation with IFNβ. We observed a large reduction in pSTAT1 signal following upadacitinib treatment, and a modest reduction in pSTAT1 signal following deucravacitinib treatment (Figure 4A-B). Treatment with both drugs also synergistically decreased the pSTAT1 signal but did not entirely ablate it, suggesting that IFNβ signaling still occurs even at high concentrations of deucravacitinib and upadacitinib (Figure 4A-B). We next tested whether the decrease in STAT1 signaling caused by these inhibitors is enough to prevent IFNβ from establishing a plasticity restriction. Macrophages were pretreated with deucravacitinib or upadacitinib before IFNβ stimulation before being washed with PBS, stimulated with IL-4, and stained for CD206 to measure the plasticity restriction. Compared to no drug treatment, hMDMs pretreated with either inhibitor, or both, saw a modest restoration in CD206 expression (Figure 4C). However, treatment with both deucravacitinib and upadacitinib at the 100 times their reported IC50 was not enough to fully rescue response to IL-4 through CD206 expression. We hypothesize that residual STAT1 phosphorylation was preventing full restoration of the naive IL-4 response.

**Figure 4:**
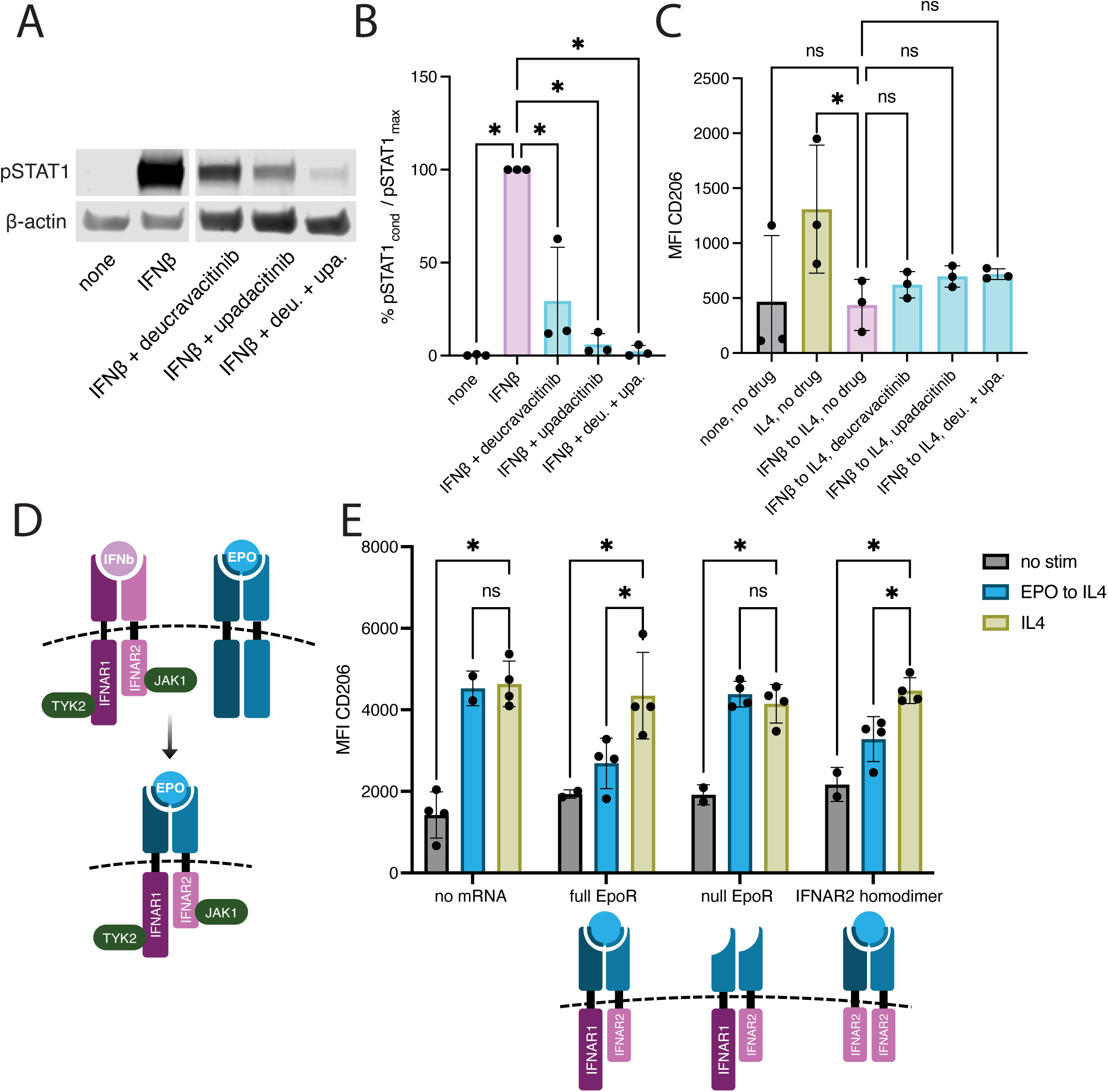
IFNAR2/JAK1 signaling is sufficient to drive plasticity restriction with respect to IL-4 **A-B)** Western blot of pSTAT1 30 minutes after IFNβ stimulation +/- 1 hour pretreatment with deucravacitinib / upadacinitib at 100X each respective IC50 (deu. = 100 nM, upa. = 4.3 μM). **A)** Representative blot. **B)** pSTAT1 signal divided by β-actin loading control signal in each well, then normalized to minimum and maximum pSTAT1 signal for each replicate. N = 3 PBMC donors stimulated on 3 separate days. One-way ANOVA performed with Dunnett correction, all conditions compared to IFNβ with no drug pretreatment. **C)** CD206 MFI above isotype for n = 3 donors on three separate days. Drug pretreatment was done for 1 hour at 100X each respective IC50 before IFNβ stimulation. One-way ANOVA performed with Dunnett correction, all conditions compared to IFNβ → IL-4, no drug condition. **D)** Schematic of EpoR-IFNAR chimeric receptor. Each respective EpoR intracellular domain is swapped for IFNAR1/IFNAR2 intracellular domain. Asymmetrical EpoR ensures that IFNAR1/2 heterodimers form upon engagement with EPO. **E)** CD206 MFI above isotype with n = 2-4 PBMC donors on three separate days, schematics of each type of receptor shown below. EPO was given at 10 ng/mL. Two-way ANOVA performed with Dunnett correction, all conditions within each group compared to each respective IL-4 stimulated condition.

To test components of the IFNβ signaling cascade individually, we utilized an established chimeric receptor system that uses the erythropoietin (EPO) extracellular domain fused to the IFNAR intracellular domain^28^, allowing us to activate IFNAR intracellular signaling specifically and independently of endogenous IFNβ receptors (Figure 4D). We delivered chimeric receptor constructs to primary human macrophages using modified nucleoside mRNAs encoding each side of the IFNAR heterodimer separated by a P2A ribosome skipping motif. Macrophages lacked the EPO receptor (EpoR) and did not respond to EPO without delivery of the chimeric receptor (Supplementary Figure 6A-B). After delivery, we pretreated macrophages with 10 ng/mL of EPO for 24 hours, then washed and replaced with 10 ng/mL of IL-4 for an additional 24 hours. We observed that EPO → IL-4 treatment recapitulated the CD206 plasticity defect seen with IFNβ → IL-4 treatment, which was dependent on presence of the chimeric receptor (Figure 4E). In addition to the fully functional EpoR-IFNAR construct, we created a null EpoR-IFNAR construct that lacks the one side of the heterodimerization site, and therefore will not signal upon contact with EPO. Additionally, we created an EpoR-IFNAR2 homodimer construct that lacks IFNAR1 and therefore only signals with JAK1 signaling. An EpoR-IFNAR1 construct was not made due to the inability of IFNAR1 to form homodimeric complexes^28,29^. As expected for the signaling-incompetent construct, macrophages expressing the null EpoR-IFNAR construct did not exhibit a plasticity restriction when treated with EPO → IL-4 (Figure 4E). Macrophages expressing the EpoR-IFNAR2 homodimer were plasticity restricted in response to IL-4, with lower expression of CD206 with EPO pretreatment. Overall, these data show that IFNAR2-mediated signaling alone is sufficient to drive a plasticity restriction, independent of IFNAR1/TYK2 signaling or IFNβ itself.

### ESX-1-dependent IFN**β** secretion during Mtb infection restricts subsequent IL-4 transcriptional responses

Having established that exogenous IFNβ restricts subsequent IL-4 responses, we next asked whether endogenously produced IFNβ during bacterial infection could establish the same plasticity restriction in a physiologically relevant context. *Mycobacterium tuberculosis* (Mtb) is a useful model due to its persistence in the macrophage phagosome^30^ and the well-characterized type I interferon response it causes during infection^31–33^. In the macrophage phagosome, Mtb secretes effectors through its type VII secretion system ESX-1^34^ which has been shown to increase cGAS-STING signaling and promote expression of IFNβ^33,35^. To test if our observations of plasticity restrictions induced by exogenous IFNβ were recapitulated by endogenous IFNβ expression caused by bacterial infection, we infected macrophages with Mtb and stimulated them with IL-4 24 hours later (Figure 5A). In addition to WT Mtb, we also infected macrophages with EccCa1:Tn Mtb, a transposon mutant in the ESX-1 machinery that abolishes secretion activity^36^, to separately test the effect of ESX-1 on macrophage transcriptional responses. We observed that a subset (29/72) of IL-4-induced genes are plasticity restricted when macrophages were infected with WT Mtb, but these genes were responsive to IL-4 when macrophages were infected with EccCa1:Tn Mtb (Figure 5B). Many of the restricted genes served functions related to cell adhesion and cytoskeletal remodeling (*TACSTD2*, *PPFIBP1*, *BIN2*, *PLXNA2*, *ADAM12*), as well as chemokines (*CCL24*, *CCL22*) and intercellular signaling (*IL1RAP*, *CD1C*, *CD1B*, *FLT1*, *MRC1*). Additionally, we observe 28 IL-4 DEGs that were plasticity restricted with exogenous IFNβ treatment as well as WT Mtb infection (Supplementary Figure 8A). These results extend our findings with recombinant IFNβ to an infection context, demonstrating that endogenously produced IFNβ is sufficient to restrict IL-4-induced gene expression programs. Furthermore, we observed an increase of IFNβ in the supernatant of macrophages infected with WT Mtb but not EccCa1:Tn Mtb relative to uninfected, confirming that ESX-1 activity during infection is driving IFNβ secretion (Figure 5C). Comparing IFNβ-induced genes with Mtb-induced genes, we observed 334 DEGs in common (Figure 5D), suggesting overlap in gene expression programs upregulated by these stimuli. To determine whether the overall transcriptional state of Mtb-infected macrophages resembles IFNβ-stimulated macrophages, we applied the PLS-DA model trained on single-cytokine stimulations to the infection dataset. We observed an increase in IFNβ prediction score for WT Mtb infected conditions relative to uninfected macrophages, as well as a slight decrease in IL-4 prediction score for WT Mtb → IL-4 polarized macrophages relative to uninfected → IL-4 (Supplementary Figure 8B). Together, these data demonstrate that infection-induced IFNβ is sufficient to restrict macrophage transcriptional plasticity, suggesting that the history of pathogen exposure can fundamentally shape the macrophage response to subsequent immunological signals.

**Figure 5:**
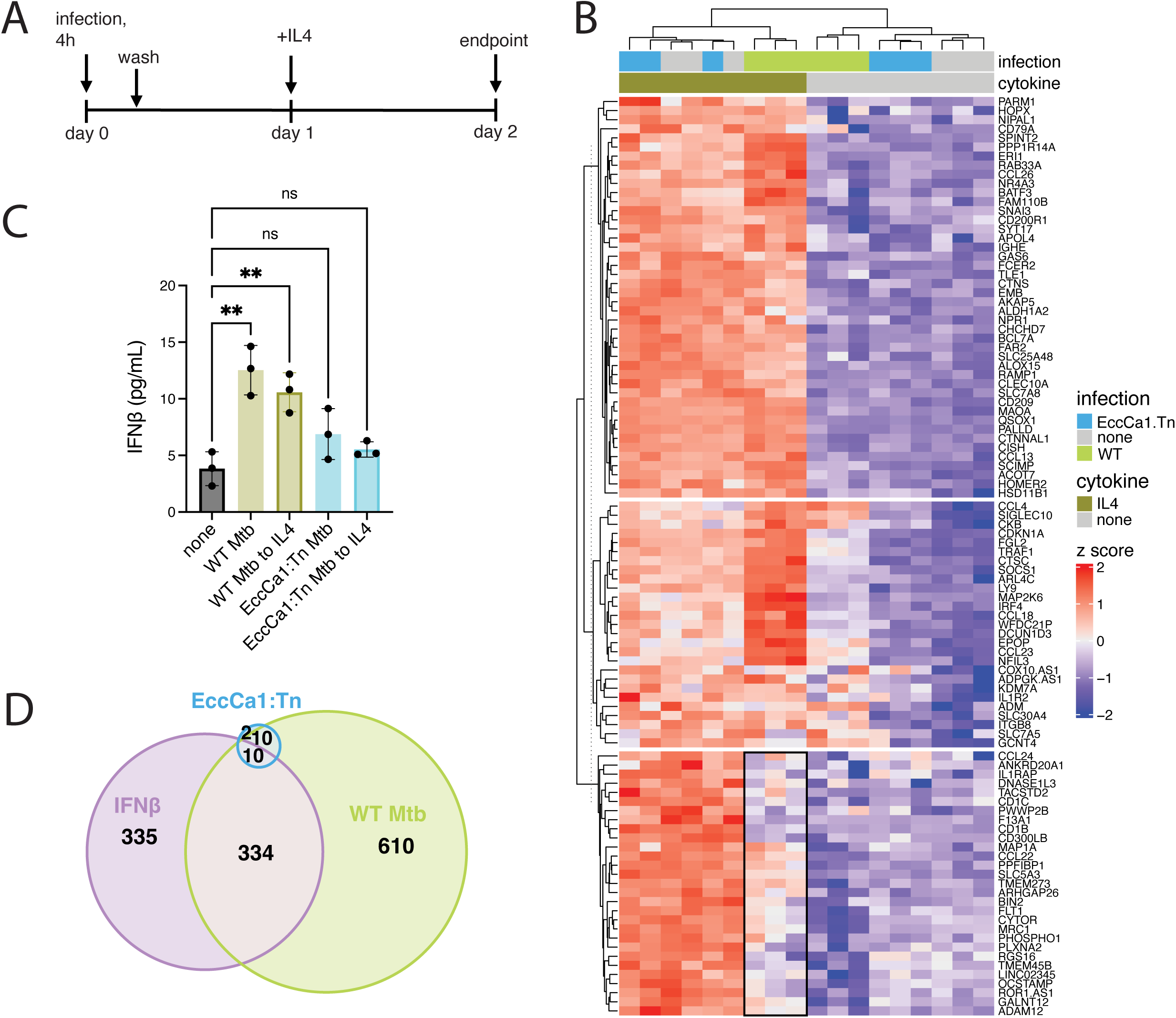
Mtb infection induces type I interferon signaling that influences subsequent IL-4 response **A)** Schematic of experimental set-up. **B)** Normalized expression of IL-4 DEGs compared to no infection / stimulation condition. K means clustered (k=3), black box around context dependent IL-4 DEGs. **C)** IFNβ ELISA from Mtb-macrophage infection media, taken from the same experiment as RNAseq. One-way ANOVA performed with Dunnett correction, all samples compared to no infection / stimulation condition. **D)** DEGs over no infection / stimulation condition, overlap proportional according to shared DEG identity. RNAseq/ELISA for **B**-**D** performed on N = 3 PBMC donors with 2 technical replicates performed on separate days.

## Discussion

Macrophage transcriptional responses to cytokine stimulation are not determined by current signals alone, but are shaped by prior activation history in a stimulus-specific and asymmetric manner. We utilized a panel of cytokines that engage STAT1, −3, and −6 as well as NF*κ*B and SMAD to analyze cytokine responses across fundamentally different transcriptional programs. We found that macrophages primed with IFNβ were especially restricted in their responses to the rest of the cytokines in the panel, both in terms of the differentially expressed genes (DEGs) associated with each cytokine as well as through cytokine prediction using partial least squares discriminant analysis (PLS-DA). These findings are consistent with prior observations that show long lasting effects of interferon pre-treatment on macrophage response to IL-4^11^ as well as TLR stimuli^12^. We expand on these findings by utilizing a broader slate of cytokines and therefore downstream gene expression programs.

We found that the order of sequential cytokine stimulation affects macrophage responses asymmetrically across cytokine pairs, as observed by differences between the forward and reverse sequences of the same two cytokines. The asymmetry we observe implies a hierarchy in cytokine responses, where the control of a subset of gene expression programs is overridden by cytokine pretreatment. We observe that IFNβ affects responses to most other cytokines, while IFNγ and IL-4 have strong but less dominant effects. This could be due to positive feedback regulation present in IFNβ signaling that sustains downstream transcriptional activity^37^. This is supported by other models of macrophage plasticity which show that interferon pretreatment results in a mixture of potentiated and diminished responses to a secondary stimulus^12^. Additionally, we observed that the IFNβ-induced plasticity restriction to the macrophage IL-4 response was tunable rather than a binary switch. We observed a significant plasticity restriction in transcriptional and protein responses to IL-4 when macrophages were pretreated with 100 pg/mL of IFNβ. This concentration is physiologically relevant as it is approximately equal to levels of IFNβ seen in the plasma of influenza patients^38^, and is likely lower than local concentrations experienced by macrophages *in vivo*.

Macrophages pretreated with 1 pg/mL of IFNβ still responded to IL-4 as naive macrophages, suggesting a minimum threshold for plasticity restrictions to take effect. These observations show that signaling intensity is relevant for tuning subsequent cytokine responses. Because macrophages in different tissue locations experience different local IFN concentrations^39^, this could produce a gradient of plasticity restriction states over a given tissue.

In addition to concentration, the duration of interferon pretreatment had a small but less influential effect on subsequent responses to IL-4. Despite our observation that the majority of IFNβ-induced genes were upregulated by 6 hours of stimulation for the conditions we tested, one hour of IFNβ pretreatment still restricted macrophage responses to IL-4. IFNβ-induced transcriptional activity has been shown to peak in activity between four and eight hours^37,40^.

While this burst in transcriptional activity is essential in driving the inflammatory gene expression programs associated with a type I interferon response, we observe that intervening with IL-4 at only one hour post IFNβ is not sufficient to disrupt this process. These data suggest that IFNβ signaling events within the first hour may be sufficient to initiate plasticity restrictions that are resistant to subsequent IL-4 stimulation.

Despite their similarities in function and signaling machinery, IFNβ drove a stronger plasticity restriction for human macrophages than IFNγ, as observed by transcriptional responses to other cytokines. Although IFNβ and IFNγ have different affinities for their respective receptors^41,42^, receptor affinity alone is not sufficient to predict downstream signaling activity^43^, suggesting that other mechanisms account for their differences in plasticity restriction. We also observed that IFNγ drives a more stable epigenetic state than IFNβ in macrophages repolarized with IL-4. This discordance in plasticity phenotypes between chromatin accessibility and transcriptional responses suggests that chromatin is accessible for IL-4-induced genes but their transcription remains affected by IFNβ pretreatment. Although we did not explicitly test post-translational regulation, it is plausible that interactions between STAT6 and its targets are hindered by other epigenetic changes besides chromatin accessibility that explain altered IL-4-induced gene expression without any effect to phosphorylation of STAT6 itself. We also observe that IFNβ-induced plasticity restrictions remain many days after cytokine wash out. This contrasts with others who report that macrophages costimulated (instead of sequentially stimulated) with pro- and anti-inflammatory cytokines revert to an IL-4-like state after several days^21^. Co-stimulation is mechanistically distinct from sequential stimulation because it prevents the establishment of a stable prior state, which may explain differences in responses. Together, this implies that more complex rules could govern macrophage state transitions over various durations, concentrations, and compositions of extracellular signaling events.

Inhibiting IFNAR signaling through TYK2 and JAK1 inhibitors did not result in complete loss of STAT1 phosphorylation during IFNβ pretreatment. Even at 100x their reported IC50s, dual inhibition was insufficient to fully rescue upregulation of CD206 following subsequent IL-4 stimulation. This could be due to residual activity of the kinases themselves or from signaling pathways that are independent of TYK2/JAK1 phosphorylation such as PI3K activity^23^. We additionally used a synthetic IFNAR receptor system to separately test components of the IFNβ signaling cascade after receptor engagement, and found that the dimerization of IFNAR2 component was sufficient to drive a plasticity restriction in response to IL-4 without using IFNβ itself. This implicates JAK1 phosphorylation specifically, although it remains unclear if this is due to STAT1-independent activity of JAK1, or if STAT1 phosphorylation is still necessary to maintain the plasticity restriction. Due to the positive feedback that is present in IFNAR signaling37, it is plausible that plasticity restrictions can arise even through low-level kinase activation as long as there is sufficient activity to trigger these regulatory mechanisms.

Infection with *Mycobacterium tuberculosis* (Mtb) resulted in a pathogen-induced plasticity restriction of a subset of IL-4-induced genes that overlapped with exogenous IFNβ-restricted genes, and was dependent on the activity of Mtb’s type VII secretion system ESX-1. Induction of a type I interferon response in macrophages infected with Mtb is generally considered to be detrimental to bacterial control^44,45^. We also observe that this type I interferon response induces a similar plasticity restriction to exogenous IFNβ, which suggests that Mtb-induced type I interferon signaling early in infection could have long-lasting effects in the TB granuloma. This has been shown to result in disrupted IFNγ macrophage responses in the lung^31^ as well as decreased cell viability *in vitro*^32^. Taken together, ESX-1-dependent IFNβ production early in infection could constrain macrophage responsiveness to subsequent immune signals, potentially contributing to the dysregulated inflammation associated with poor TB outcomes. Broadly, type I interferons have been implicated in a variety of bacterial infection contexts despite their association with antiviral responses^46^. For example, type I interferon signaling in macrophages infected with *Listeria monocytogenes* results in the downregulation of the IFNγ receptor (IFNGR), which in turn increases susceptibility to infection and spread^47^. This IFNβ-induced disruption in subsequent cytokine response mirrors our observations with Mtb infection, although we do not directly observe IL-4 receptor downregulation. Conversely, *Shigella sonnei* infection blocks interferon signaling through effectors secreted through its type III secretion system^48^. Together, our findings extend the known consequences of type I interferon signaling during bacterial infection to include durable restrictions on macrophage transcriptional plasticity that are mediated by the activity of a bacterial secretion system.

## Limitations of the study

While bulk transcriptomic data provides a useful framework for analyzing global macrophage state, plasticity restrictions and their mechanisms span modes of regulation that are not well captured by changes in mRNA at a cell population level. For example, macrophage state is highly intertwined with metabolic state; IFN-stimulated macrophages upregulate glycolysis and IL-4-stimulated macrophages upregulate oxidative phosphorylation^49^. This metabolic switch has been implicated in restricting macrophage cytokine responses^11^, as well as responses to bacterial infection^50^. The plasticity restriction we observe in CD206 protein expression is not explained by *MRC1* transcript, which implies post-transcriptional regulation. Proteomic analysis following sequential cytokine stimulation would directly address the discrepancies we observe between transcript and protein expression that occur after IFNβ pretreatment. Interferon-induced epigenetic changes that are not captured by changes in chromatin accessibility could play a role in limiting macrophage plasticity long term. Other targets of interferon stimulation that are STAT1-independent, such as P13K, were not explored but could be important in contributing to changes in macrophage signaling intracellularly. While monocyte-derived macrophages are a useful *in vitro* model of primary human cell biology, tissue resident macrophages likely exhibit differences in plasticity depending on tissue type.

## Materials and methods

### Primary macrophage cell culture

For primary human macrophages, buffy coats from healthy human donors at Massachusetts General Hospital were collected. PBMCs were isolated via centrifugation in Ficoll Paque Plus (Millipore Sigma, cat #GE17-1440-03). Monocytes were then subsequently isolated using CD14+ positive selection (Stemcell, cat #17858) according to manufacturer’s instructions, and differentiated for 6 days in phenol-free RPMI (Thermo Fisher, cat #11835030), 10% FBS (Thermo Fisher, cat #10082147), 2 mM L-glutamine (Sigma, cat #G7513), 10 mM HEPES (Corning, cat #25-060-CI), and 25 ng/mL M-CSF (Biolegend, cat #574804) on ultra-low attachment plates (Corning, cat #3814). Macrophages were detached with 4 mM EDTA (Corning, cat #46-042-CI) for 15 minutes, and replated in RPMI, 10% FBS, 2 mM L-glutamine, and 10 mM HEPES without M-CSF. For the cytokine stimulations, IFNβ (PeproTech cat #300-02BC), IFNγ (Biolegend, cat #570204), IL-1β (Biolegend, cat #579404), IL-4 (Biolegend, cat #574004), IL-10 (Biolegend, cat #571004), TGFβ (PeproTech, cat #100-21C), and erythropoietin (Sigma, cat #E5546) were diluted in supplemented RPMI without M-CSF. For the repolarization conditions, three DPBS (Corning, cat #21-031-CV) washes were performed between each cytokine.

### Bulk RNA sequencing library prep

Macrophages were lysed at a concentration of 1,000 cells per µL in RLT plus buffer (Qiagen cat #1053393) with 1% β-mercaptoethanol (Sigma, cat #M31-48) and frozen in 25 µL aliquots at −80 ℃ until use. Bulk RNA library prep was performed following a modified Primeseq protocol51. RNAclean Ampure XP beads (Beckman, cat #A63881) were added to lysates at a 2:1 ratio for 5 minutes, washed twice with 80% ethanol, and eluted in 4 µL nuclease-free water. To each sample, 1 µL of barcode oligos and 5 µL of reverse transcription master mix was added, and the remaining library preparation was performed as described in the original protocol. Libraries were sequenced with a 75 cycle high-output kit on a NextSeq500 or a 100 cycle P3 XLEAP-SBS kit on a NextSeq2000.

### Gene expression analysis

Base call files were converted to .fastq using bcl2fastq with adapter trimming. Data was aligned and filtered using the zUMIs pipeline^52^. Differential gene expression analysis was performed with DESeq2^53^, with log_2_ fold changes shrunk using apeGLM^54^. Differential expression was determined with an adjusted p value < 0.05 and the absolute value of shrunk log_2_ fold change > 1. PLS-DA was performed with the scikit-learn PLSRegression function on a subset of genes that were detected in all experiments to allow for use across datasets (9,889 genes). The model was trained on all conditions with a single identity of cytokine (“none → cytokine X”, “cytokine X → none”, “cytokine X → cytokine X”) for all donors. Regression was performed using a one-hot encoded response matrix with 7 columns (the 6 cytokines plus no stimulation), where each sample was assigned a value of 1 for the column corresponding to that cytokine and a 0 for every other column.

### Flow cytometry

Macrophages were detached using 4 mM EDTA for 15 minutes, then spun at 500xg and resuspended in 1:1000 LIVE/DEAD Fixable Near-IR dye (Thermo Fisher, cat #L10119) in DPBS for 10 minutes at room temperature. Cells were spun again and resuspended in 1:250 Human TruStain FcX (Biolegend, cat #422302) diluted with FACS buffer (DPBS supplemented with 2% FBS and 4 mM EDTA) and incubated at room temperature for 10 minutes. Cells were then stained with primary conjugated antibodies (anti-CD206: Biolegend cat #321105, anti-erythropoietin receptor: Biotechne cat #FAB307P, mouse IgG1 isotype: Biolegend cat #400111, mouse IgG2 isotype: Biolegend cat #400211) diluted 1:250 in FACS buffer, added on top of Fc block, and incubated at room temperature for 15 minutes. Cells were spun and then fixed for 15 minutes at room temperature with 4% paraformaldehyde (Santa Cruz, cat #30525-89-4), washed twice, and resuspended in FACS buffer before being analyzed on a BD FACSymphony cytometer.

### Western blot

Cells were lysed in RIPA with Triton X-100 (Thermo Fisher, cat #J62725) with 1:100 Halt protease inhibitor cocktail (Thermo Fisher, cat #78429). For samples that were stained for phosphorylated proteins, phosphatase inhibitor cocktail was added 1:100 (Thermo Fisher, cat #J63907). For CD206 western blots, lysis buffer was supplemented with 1% SDS (Thermo Fisher, cat #15553027) to improve dissolution of the protein from the membrane. The 1% SDS samples were subsequently treated with benzonase (Sigma, cat #70664-3) 1:1,000 for 30 minutes at room temperature to eliminate excess viscosity. Lysates were clarified by centrifugation at 10,000xg for 10 minutes before transferring the supernatant to a new tube.

Clarified lysates were quantified using Pierce 660nm Protein Assay Reagent (Thermo Fisher, cat #22660) using Ionic Detergent Compatibility Reagent (Thermo Fisher, cat #22663) according to the manufacturer’s instructions. Samples were boiled at 95 ℃ for 5 minutes in Laemmli SDS sample buffer and reducing agent (Thermo Fisher, cat #J61337), loaded onto a 4-12% Bis-Tris gel (Thermo Fisher, cat #NW04125), and transferred to a nitrocellulose membrane using an iBlot2 dry transfer system (Invitrogen). Membranes were blocked for 1 hour at room temperature in TBS blocking buffer (Licor, cat #927-60001), then stained shaking overnight at 4℃ in 1:1,000 primary antibody (Cell Signal Technology: anti-phosphoSTAT1 cat #9167, anti-STAT6 cat #5397, anti-phosphoSTAT6 cat #56554, Thermo Fisher: anti-CD206 cat #MA5-28581, Biolegend: anti-β-actin cat #664802). Membranes were washed three times in TBS-T (Thermo Fisher, cat #J60764), then incubated with 1:10,000 secondary antibody (Licor: goat anti-rabbit 800CW cat #926-32211, goat anti-mouse 800CW cat #926-32210, goat anti-rat 680RD cat #925-68076) for 1 hour at room temperature, then washed three more times in TBS-T before imaging on a Licor Odyssey CLx imager.

### mRNA synthesis and delivery

EpoR-IFNAR component plasmids were received from the Alexander Ploss Lab. Each construct was assembled via Gibson reaction into an *in vitro* transcription (IVT) backbone which contains 5’ and 3’ UTRs and a polyA tail. Assemblies were transformed into DH5α competent E. coli (New England Biolabs, cat #C2987H), grown in LB broth (Sigma, cat #L3522) and miniprepped (Omega Bio-Tek, cat #D6942-03). 1-2 µg of plasmid was linearized at the 3’ end of the polyA tail and purified using QIAquick PCR purification kit (Qiagen, cat #28104). The template DNA was then transcribed using HiScribe T7 mRNA Kit with CleanCap Reagent (New England Biolabs, cat #E2080S) using methoxyuridine (Trilink, cat #N-1093-10) and methylcytidine (Trilink, cat #N-1014-10) for efficient delivery in macrophages. mRNA was purified using Monarch RNA cleanup kit (New England Biolabs, cat #T2040L).

To deliver mRNA to macrophages, 7.5 µL of Lipofectamine MessengerMAX transfection reagent (Thermo Fisher, cat #LMRNA008) was added to 125 µL Opti-MEM (Thermo Fisher, cat #31985062) per million cells, and incubated for 10 minutes at room temperature. 3 pmol of mRNA per million cells was diluted with an additional 125 µL Opti-MEM, then added to the lipofectamine/Opti-MEM mixture and incubated for 5 minutes at room temperature. The mixture was added to macrophages without media removal before a 24 hour incubation.

### Luminex

For the signaling pathway validation experiment, macrophages were stimulated with IFNβ, IFNγ, IL-1β, IL-4, IL-10, or TGFβ for 30 minutes before lysis and normalization of protein concentration. Lysates were arrayed in 384-well plates for efficient use of materials, and incubated with beads from the MILLIPLEX STAT and TGFβ signaling panels (Sigma: STAT kit cat #48-610MAG, TGFβ kit cat #48-614MAG) shaking overnight at 4 ℃. Washes and antibody incubations were performed according to the manufacturer’s instructions. Samples were analyzed on a FlexMap 3D Luminex plate reader.

### Single-cell ATACseq library prep and analysis

Nuclei were isolated from 1 million macrophages per condition using lysis buffer: nuclease-free water with 10 mM Tris-HCl (Thermo Fisher, cat #15567027), 10 mM NaCl (Thermo Fisher, cat #AM9760G), 3 mM MgCl_2_ (Sigma, cat #7786-30-3), 0.1% Tween-20 (Sigma, cat #9005-64-5), 0.1% IGEPAL CA-630 (Sigma, cat #9002-93-1), 0.01% Digitonin (Promega, cat #G9441), and 1% BSA (Thermo Fisher cat #AM2616). Samples were loaded onto a Chromium Controller (10X genomics) targeting 8,000 nuclei / sample. Libraries were prepared according to the 10X Genomics ATACseq protocol (CG000496 Rev C). Gene expression and antibody libraries were sequenced with a 150 cycle high-output kit on a NextSeq500. Data was demultiplexed, aligned, and quantified using cellranger-atac. Transcription factor motif enrichment was quantified using chromVAR^55^.

### Mycobacterial culture and infection

*Mycobacterium tuberculosis (*Mtb) H37Rv was cultured with Difco Middlebrook 7H9 (VWR, cat #90003-876) media with 10% oleic acid dextrose catalase (OADC) (Fisher Scientific, cat #B12351), 0.2% glycerol (Sigma, cat #G6279), and 0.05% Tween-80 (Sigma, cat #P1754) with kanamycin (Sigma, cat #K0254). Mtb strains used were GFP+ H37Rv (KanR), and GFP+ EccCa1:Tn H37Rv (KanR). Bacteria were grown to mid-log phase, washed with PBS, and resuspended in supplemented RPMI media described above. Bacteria were then spun at 500 rpm to pellet larger clumps. hMDMs were infected at a multiplicity of infection (MOI) of 2 for 4 hours. Afterwards, hMDMs were washed with PBS to remove extracellular bacteria and cultured for 24 hours in plain media or IFNβ supplemented media. Cells were again washed with PBS and cultured in plain or IL-4 supplemented media for an additional 24 hours.

Supernatants were saved, then cells were lysed with RLT plus 1% β-mercaptoethanol. Supernatants and lysates were sterile filtered twice using 0.2 µm filter plates and stored at −80℃ until use.

### ELISA

IFNβ ELISA was performed using ELISA MAX Deluxe human IFNβ kit (Biolegend, cat #449504) with ELISA plates (Biolegend, cat #430604) according to the manufacturer’s instructions.

Supernatants were diluted using Assay Diluent B in the ELISA kit prior to addition to the assay.

## Acknowledgements

We thank the entire Bryson lab for their feedback and scientific discussions. Diane Ballestas provided administrative support. Ryan Milligan and Iris Abrahantes-Morales processed buffy coats. Flow cytometry was performed at the Ragon Institute flow cytometry core managed by Michael Waring, Nathalie Bonheur, and Kat Folz-Donahue. Mtb infections were performed at the Ragon Institute BSL3 facility which is managed by Amy Barczak, Yong Xie, Julie Boucau and Eliane Shwiari. The EccCa1:Tn H37Rv GFP and WT H37Rv GFP Mtb strains were provided by Amy Barczak and Charul Jani. Sequencing libraries are quality checked by student workers at the MIT BioMicrocenter managed by Stuart Levine and supported in part by the Koch Institute Support (core) Grant P30-CA14051 from the National Cancer Institute. Alexander Ploss and Brigitte Heller provided the EpoR-IFNAR constructs. This work was supported by the National Institute of Allergy and Infectious Disease (R01AI022553, R01AI184666) and the U.S. Department of Defense (W911NF-19-D-0001).

---

**Supplementary Figure 1: Expected downstream phosphorylation cascade responds to each cytokine at a range of concentrations**. N = 4 PBMC donors x 2 technical replicates performed on different days.

**Supplementary Figure 2: DEGs in repolarized conditions are defined by presence in each single cytokine condition. A)** Single cytokine DEGs are defined by differential expression with cytokine treatment on the same day as the repolarized condition. In this example, IFNβ DEGs are determined by “IFNβ → none” compared to “none → none” and IL-4 DEGs are determined by “none → IL-4” compared to “none → none”. Genes that were DE for both cytokines were excluded. **B)** Emergent DEGs for each repolarization condition, where emergent DEGs are defined as genes not differentially expressed in either “cytokine A → none” or “cytokine B → none”.

**Supplementary Figure 3: IFNβ, IL-4, and IFNγ drive variance for PC1, PC2, and PC3, respectively A)** Principal component analysis split by cytokine one identity for PC1 and PC2 in violins plots. Circle color corresponds to cytokine 2 identity. **B)** Top 30 loadings from PCA for components 1-3.

**Supplementary Figure 4:** PLS-DA scores for all classes

**Supplementary Figure 5: CD206 protein expression is plasticity restricted. A)** Western blot of total cellular CD206 protein expression, representative of n=2. B) Induction of CD206 surface expression by IL-4 is not rescued by increasing the time between IFNβ removal and addition of IL-4, representative of n=2.

**Supplementary Figure 6: Macrophage epigenetic state is differentially stable after type I and type II interferon pretreatment A)** UMAP of single-cell ATACseq performed on nuclei isolated from macrophages polarized with 10 ng/mL of each cytokine for 24 hours or repolarized for an additional 24 hours. **B)** Pearson correlation between conditions based on transcriptional factor motif enrichment. **C-D)** Transcription factor enrichment scores from single-cell ATAC performed on repolarized macrophages. Transcription factor motifs plotted are highest variance in **C)** interferon conditions or **D)** IL-4 conditions.

**Supplementary Figure 7: EpoR-IFNAR system functions orthogonally in primary human macrophages A)** EpoR-IFNAR mRNA is efficiently delivered to primary human macrophages. No mRNA condition received lipofectamine without mRNA. Representative of n=3. **B)** EPO stimulation at 10 ng/mL results in STAT1 phosphorylation that is specific to conditions transfected with EpoR-IFNAR mRNA. LF only = lipofectamine only. EpoR = full EpoR-IFNAR construct. tSTAT1 = total STAT1. Representative of n=3.

**Supplementary Figure 8: Macrophage transcriptional response to IL-4 stimulation after Mtb infection shares features with exogenous IFNβ pretreatment A)** DEGs relative to first day stimulation only (either none, IFNβ, or WT Mtb infection). Overlap proportional according to shared DEG identity. **B)** PLS-DA prediction score for Mtb-infected macrophages. Classes with scores > 0.3 in at least one condition shown. PLS trained on subset of genes present in training dataset as well as Mtb infection dataset.

