## Supplemental Figure 1 for "Macrophage signaling history constrains subsequent transcriptional responses to cytokine stimulation"

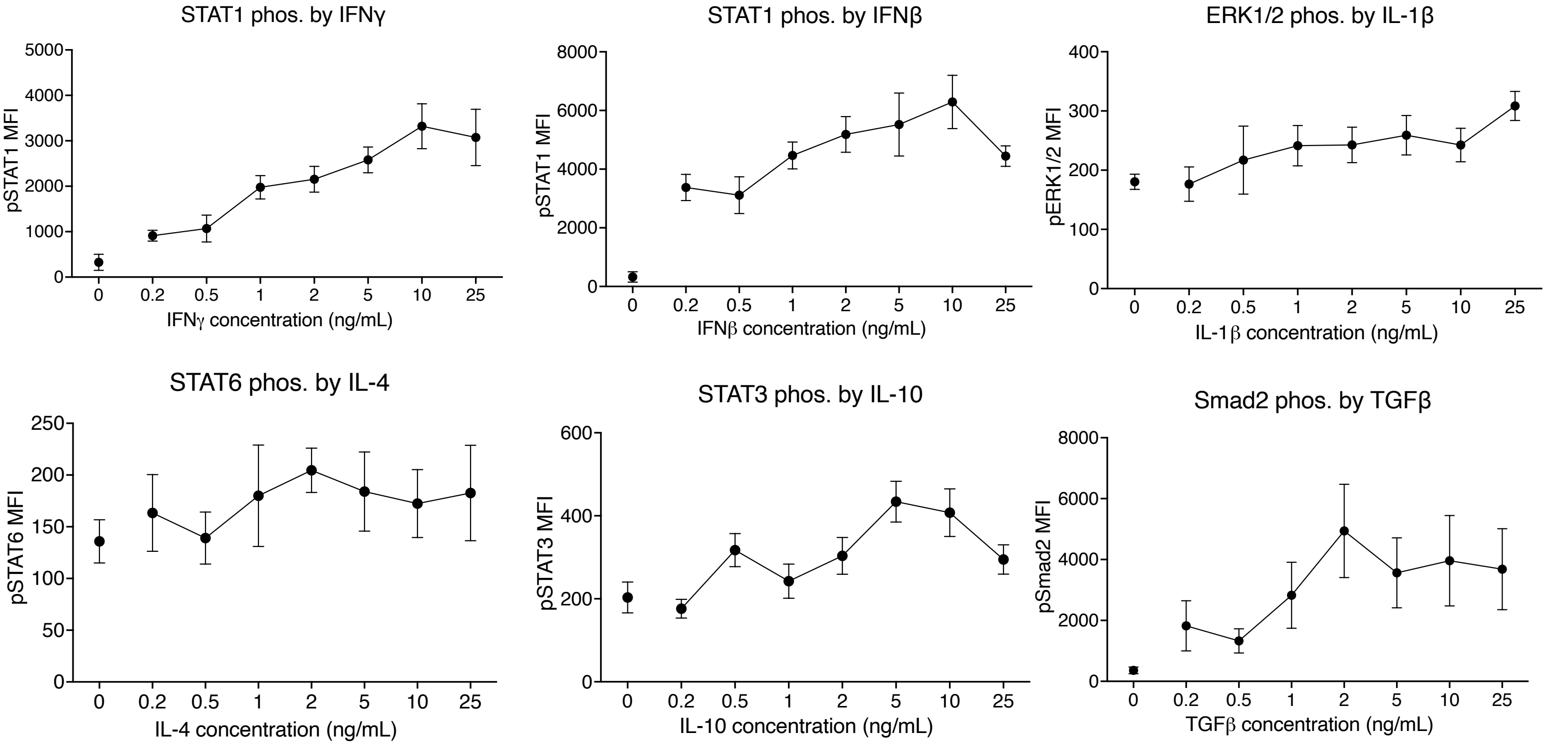

STAT1 phos. by IFNγ
pSTAT1 MFI
5000
4000
3000
2000
1000
0
0
0.2
0.5
1
2
5
10
25
IFNγ concentration (ng/mL)
STAT1 phos. by IFNβ
pSTAT1 MFI
8000
6000
4000
2000
0
IFNβ concentration (ng/mL)
ERK1/2 phos. by IL-1β
pERK1/2 MFI
400
300
200
100
0
IL-1β concentration (ng/mL)
STAT6 phos. by IL-4
pSTAT6 MFI
250
200
150
100
50
0
IL-4 concentration (ng/mL)
STAT3 phos. by IL-10
pSTAT3 MFI
600
400
200
0
IL-10 concentration (ng/mL)
Smad2 phos. by TGFβ
pSmad2 MFI
8000
6000
4000
2000
0
TGFβ concentration (ng/mL)
