## Supplemental Figure 2 for "Macrophage signaling history constrains subsequent transcriptional responses to cytokine stimulation"

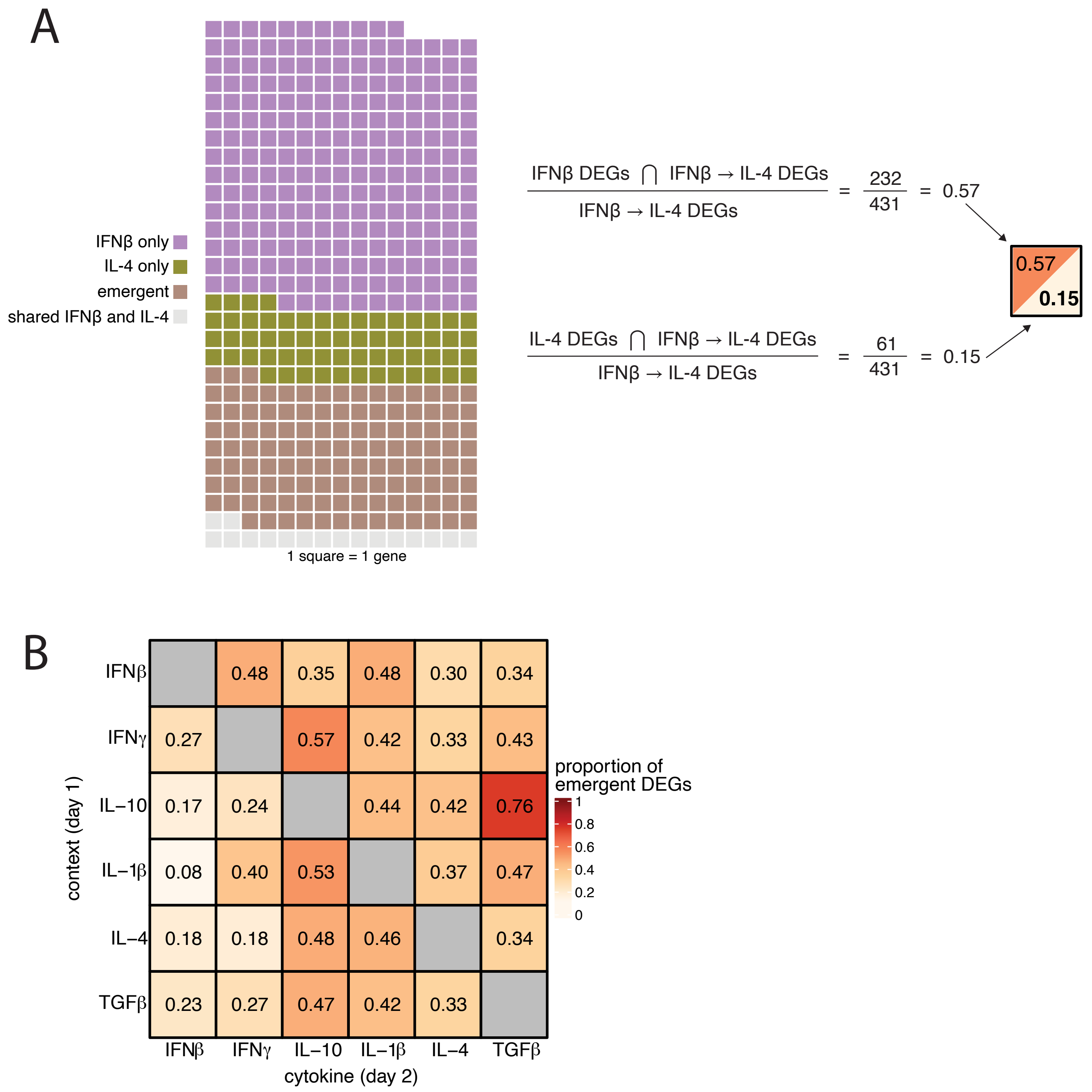
A
IFNβ only
IL-4 only
emergent
shared IFNβ and IL-4
1 square = 1 gene
IFNβ DEGs ⋂ IFNβ → IL-4 DEGs / IFNβ → IL-4 DEGs = 232 / 431 = 0.57
IL-4 DEGs ⋂ IFNβ → IL-4 DEGs / IFNβ → IL-4 DEGs = 61 / 431 = 0.15
0.57
0.15
B
context (day 1)
cytokine (day 2)
IFNβ IFNγ IL−10 IL−1β IL−4 TGFβ
IFNβ: 0.48 0.35 0.48 0.30 0.34
IFNγ: 0.27 0.57 0.42 0.33 0.43
IL−10: 0.17 0.24 0.44 0.42 0.76
IL−1β: 0.08 0.40 0.53 0.37 0.47
IL−4: 0.18 0.18 0.48 0.46 0.34
TGFβ: 0.23 0.27 0.47 0.42 0.33
proportion of
emergent DEGs
1
0.8
0.6
0.4
0.2
0
