## Supplemental Figure 3 for "Macrophage signaling history constrains subsequent transcriptional responses to cytokine stimulation"

A

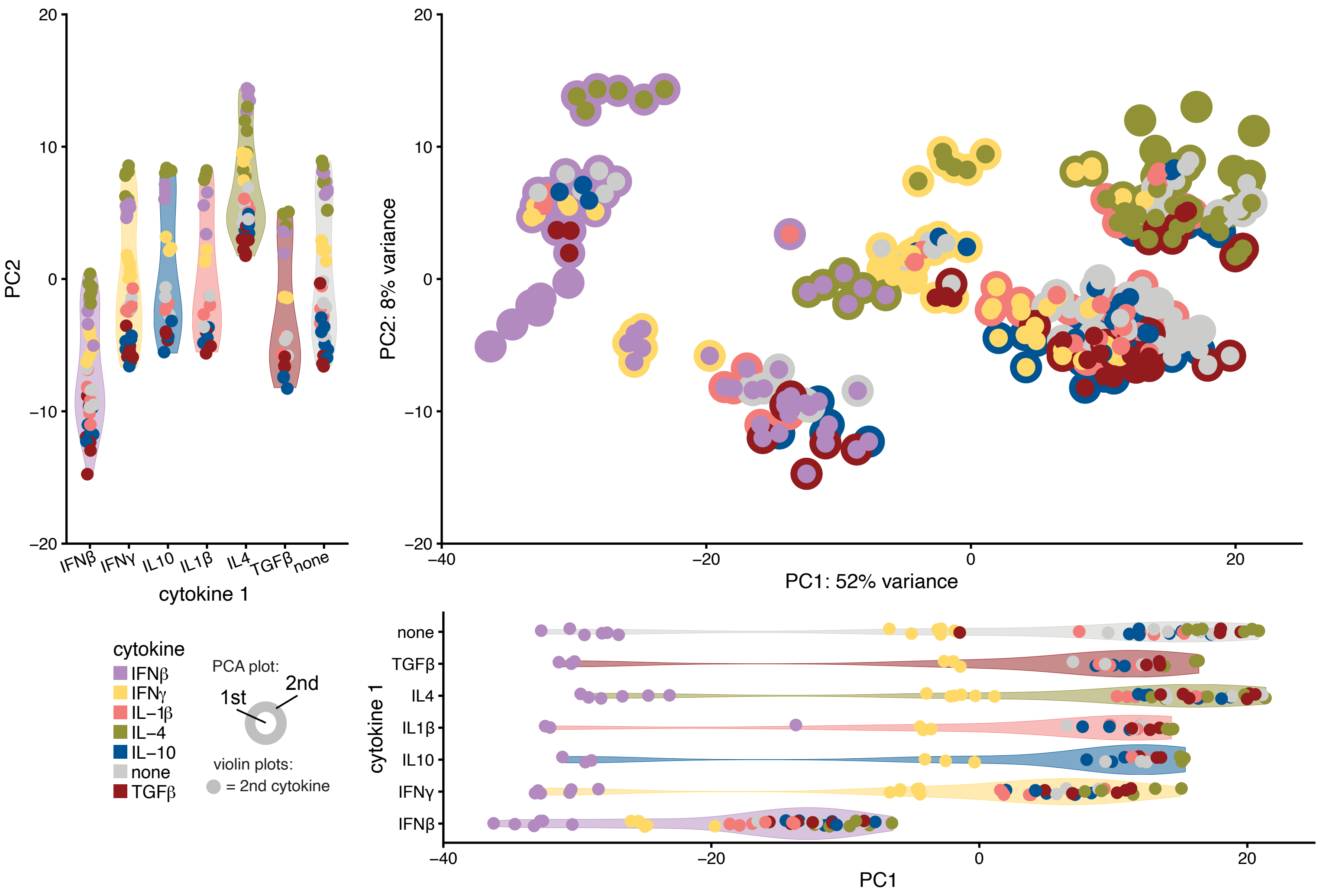
PC2
cytokine 1
IFNβ
IFNγ
IL10
IL1β
IL4
TGFβ
none
PC2: 8% variance
PC1: 52% variance
cytokine
IFNβ
IFNγ
IL−1β
IL−4
IL−10
none
TGFβ
PCA plot:
1st
2nd
violin plots:
= 2nd cytokine
cytokine 1
PC1

B

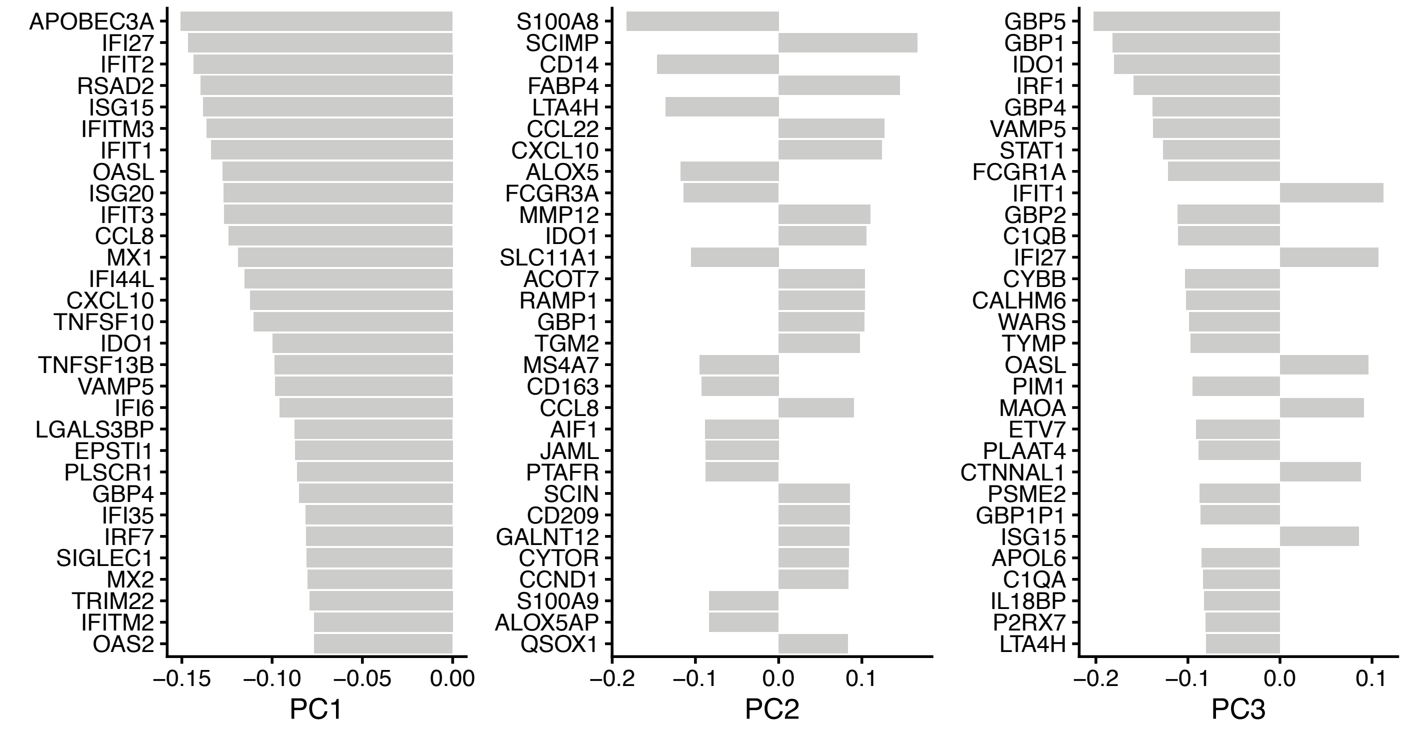
APOBEC3A
IFI27
IFIT2
RSAD2
ISG15
IFITM3
IFIT1
OASL
ISG20
IFIT3
CCL8
MX1
IFI44L
CXCL10
TNFSF10
IDO1
TNFSF13B
VAMP5
IFI6
LGALS3BP
EPSTI1
PLSCR1
GBP4
IFI35
IRF7
SIGLEC1
MX2
TRIM22
IFITM2
OAS2
PC1
S100A8
SCIMP
CD14
FABP4
LTA4H
CCL22
CXCL10
ALOX5
FCGR3A
MMP12
IDO1
SLC11A1
ACOT7
RAMP1
GBP1
TGM2
MS4A7
CD163
CCL8
AIF1
JAML
PTAFR
SCIN
CD209
GALNT12
CYTOR
CCND1
S100A9
ALOX5AP
QSOX1
PC2
GBP5
GBP1
IDO1
IRF1
GBP4
VAMP5
STAT1
FCGR1A
IFIT1
GBP2
C1QB
IFI27
CYBB
CALHM6
WARS
TYMP
OASL
PIM1
MAOA
ETV7
PLAAT4
CTNNAL1
PSME2
GBP1P1
ISG15
APOL6
C1QA
IL18BP
P2RX7
LTA4H
PC3
