## Supplemental Figure 4 for "Macrophage signaling history constrains subsequent transcriptional responses to cytokine stimulation"

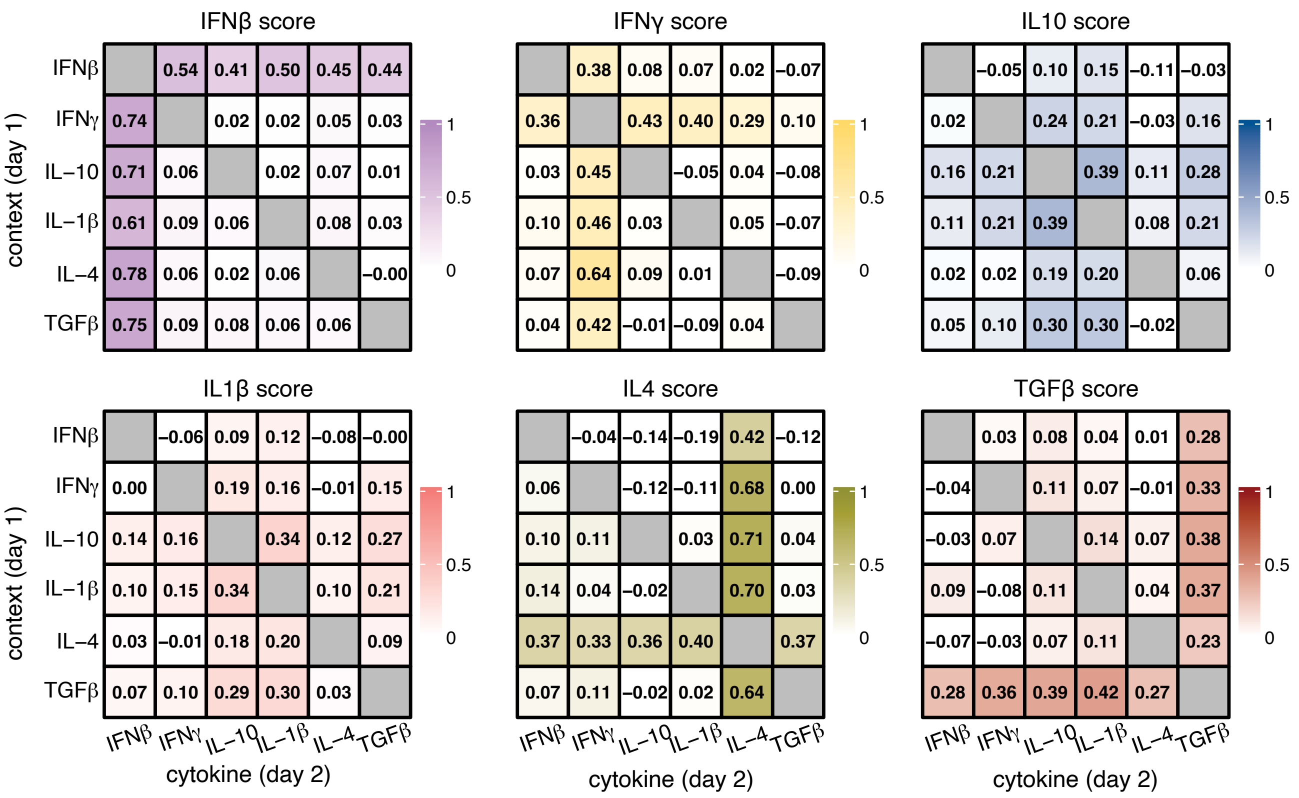
IFNβ score
IFNβ 0.54 0.41 0.50 0.45 0.44
IFNγ 0.74 0.02 0.02 0.05 0.03
IL–10 0.71 0.06 0.02 0.07 0.01
IL–1β 0.61 0.09 0.06 0.08 0.03
IL–4 0.78 0.06 0.02 0.06 –0.00
TGFβ 0.75 0.09 0.08 0.06 0.06
IFNγ score
0.38 0.08 0.07 0.02 –0.07
0.36 0.43 0.40 0.29 0.10
0.03 0.45 –0.05 0.04 –0.08
0.10 0.46 0.03 0.05 –0.07
0.07 0.64 0.09 0.01 –0.09
0.04 0.42 –0.01 –0.09 0.04
IL10 score
–0.05 0.10 0.15 –0.11 –0.03
0.02 0.24 0.21 –0.03 0.16
0.16 0.21 0.39 0.11 0.28
0.11 0.21 0.39 0.08 0.21
0.02 0.02 0.19 0.20 0.06
0.05 0.10 0.30 0.30 –0.02
IL1β score
IFNβ –0.06 0.09 0.12 –0.08 –0.00
IFNγ 0.00 0.19 0.16 –0.01 0.15
IL–10 0.14 0.16 0.34 0.12 0.27
IL–1β 0.10 0.15 0.34 0.10 0.21
IL–4 0.03 –0.01 0.18 0.20 0.09
TGFβ 0.07 0.10 0.29 0.30 0.03
IL4 score
–0.04 –0.14 –0.19 0.42 –0.12
0.06 –0.12 –0.11 0.68 0.00
0.10 0.11 0.03 0.71 0.04
0.14 0.04 –0.02 0.70 0.03
0.37 0.33 0.36 0.40 0.37
0.07 0.11 –0.02 0.02 0.64
TGFβ score
0.03 0.08 0.04 0.01 0.28
–0.04 0.11 0.07 –0.01 0.33
–0.03 0.07 0.14 0.07 0.38
0.09 –0.08 0.11 0.04 0.37
–0.07 –0.03 0.07 0.11 0.23
0.28 0.36 0.39 0.42 0.27
context (day 1)
IFNβ IFNγ IL–10 IL–1β IL–4 TGFβ
cytokine (day 2)
1
0.5
0
