## Supplemental Figure 5 for "Macrophage signaling history constrains subsequent transcriptional responses to cytokine stimulation"

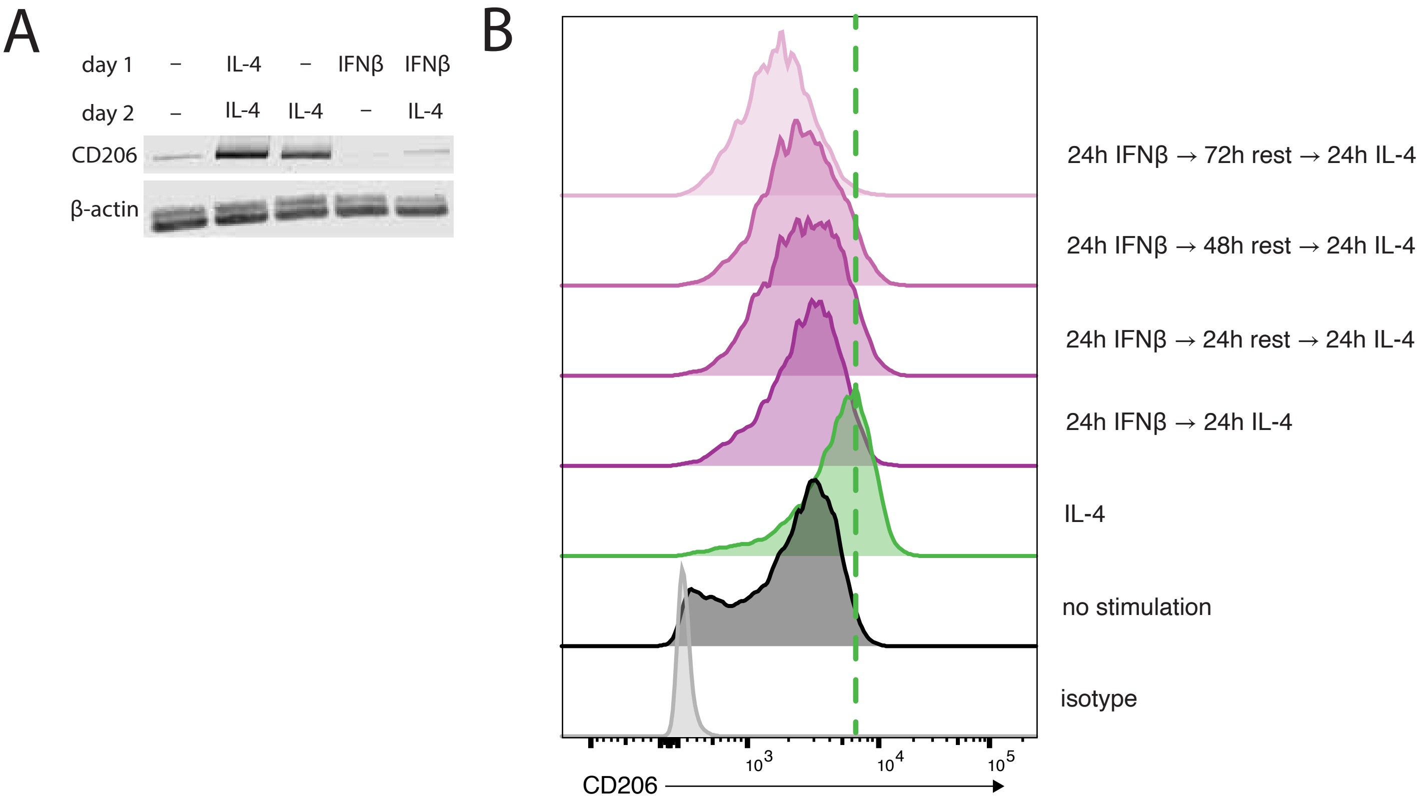
A
day 1
–
IL-4
–
IFNβ
IFNβ
day 2
–
IL-4
IL-4
–
IL-4
CD206
β-actin
B
24h IFNβ → 72h rest → 24h IL-4
24h IFNβ → 48h rest → 24h IL-4
24h IFNβ → 24h rest → 24h IL-4
24h IFNβ → 24h IL-4
IL-4
no stimulation
isotype
10^3
10^4
10^5
CD206
