## Supplemental Figure 6 for "Macrophage signaling history constrains subsequent transcriptional responses to cytokine stimulation"

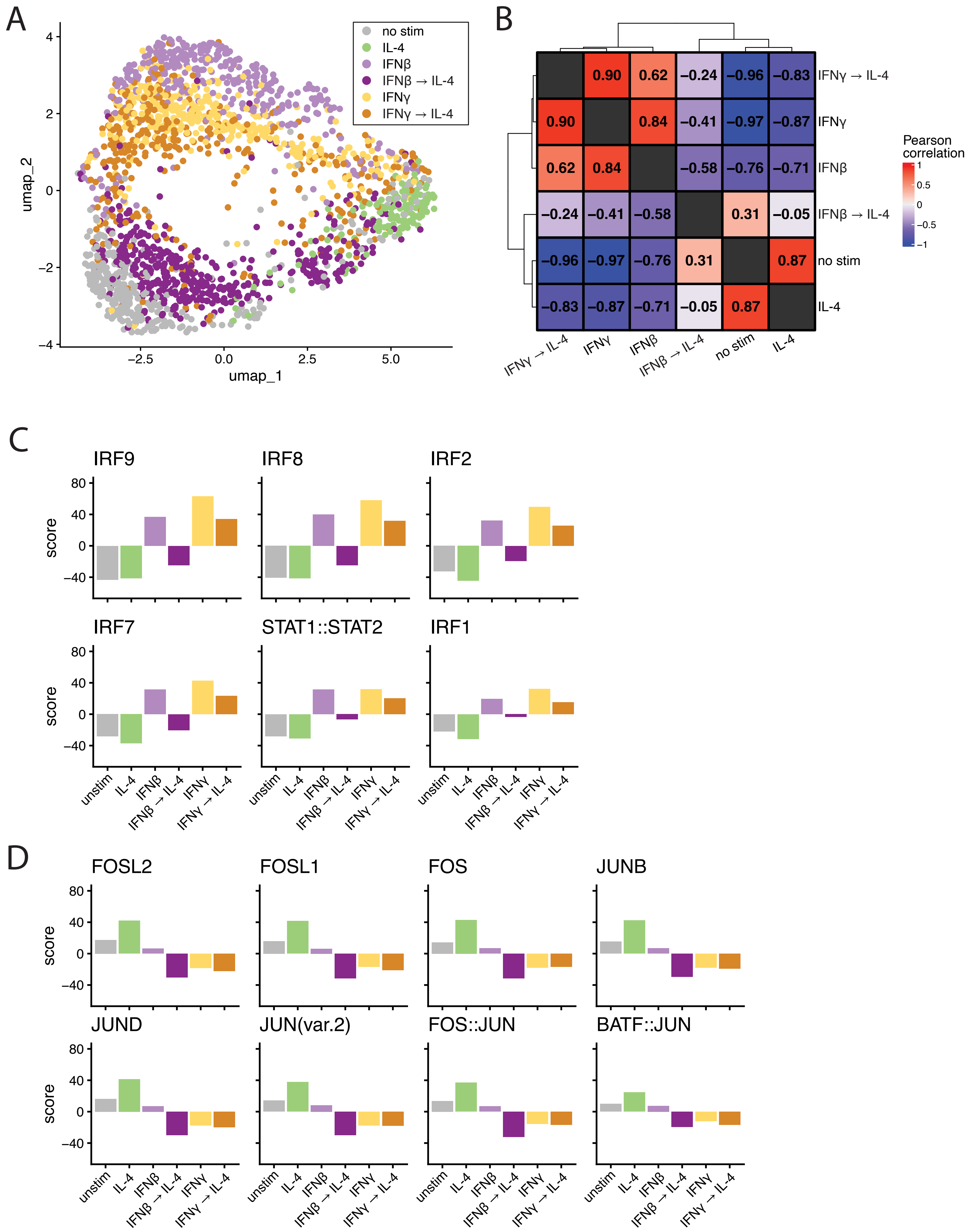

A
no stim
IL-4
IFNβ
IFNβ → IL-4
IFNγ
IFNγ → IL-4
umap_2
umap_1
B
Pearson correlation
IFNγ → IL-4
IFNγ
IFNβ
IFNβ → IL-4
no stim
IL-4
0.90 0.62 −0.24 −0.96 −0.83
0.90 0.84 −0.41 −0.97 −0.87
0.62 0.84 −0.58 −0.76 −0.71
−0.24 −0.41 −0.58 0.31 −0.05
−0.96 −0.97 −0.76 0.31 0.87
−0.83 −0.87 −0.71 −0.05 0.87
C
IRF9
IRF8
IRF2
IRF7
STAT1::STAT2
IRF1
score
unstim
IL-4
IFNβ
IFNβ → IL-4
IFNγ
IFNγ → IL-4
D
FOSL2
FOSL1
FOS
JUNB
JUND
JUN(var.2)
FOS::JUN
BATF::JUN
score
