## Supplemental Figure 7 for "Macrophage signaling history constrains subsequent transcriptional responses to cytokine stimulation"

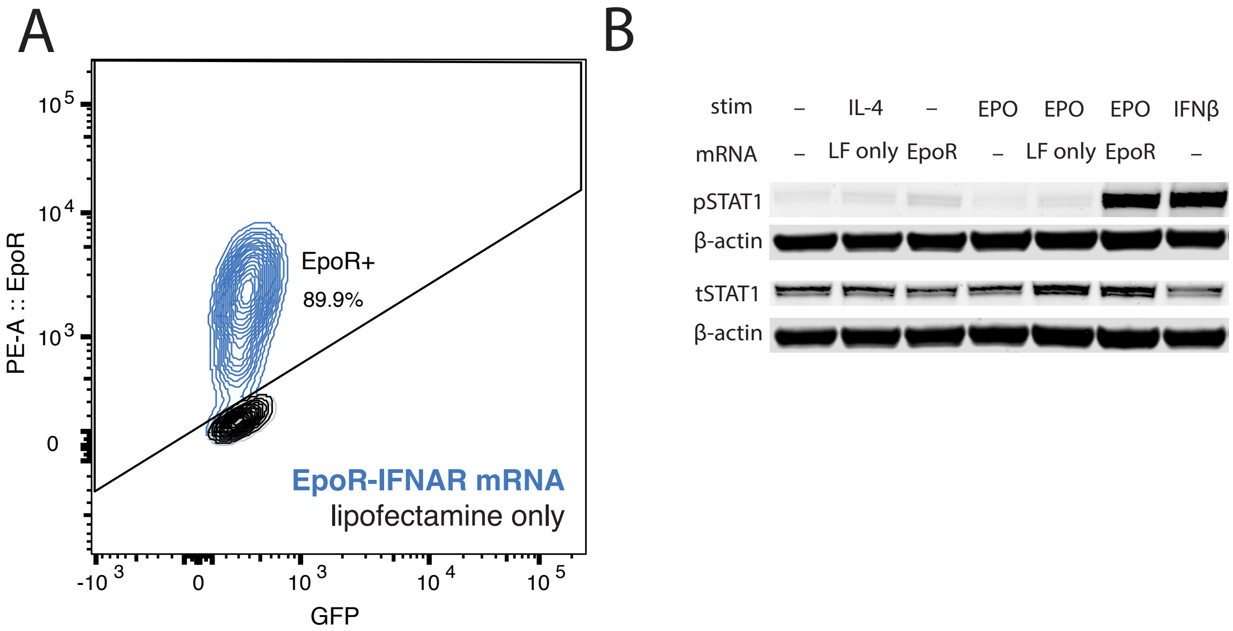
A
B
10^5
10^4
10^3
0
-10^3
PE-A :: EpoR
GFP
EpoR+
89.9%
EpoR-IFNAR mRNA
lipofectamine only
stim
mRNA
–
IL-4
EPO
IFNβ
LF only
EpoR
pSTAT1
β-actin
tSTAT1
