## Supplemental Figure 8 for "Macrophage signaling history constrains subsequent transcriptional responses to cytokine stimulation"

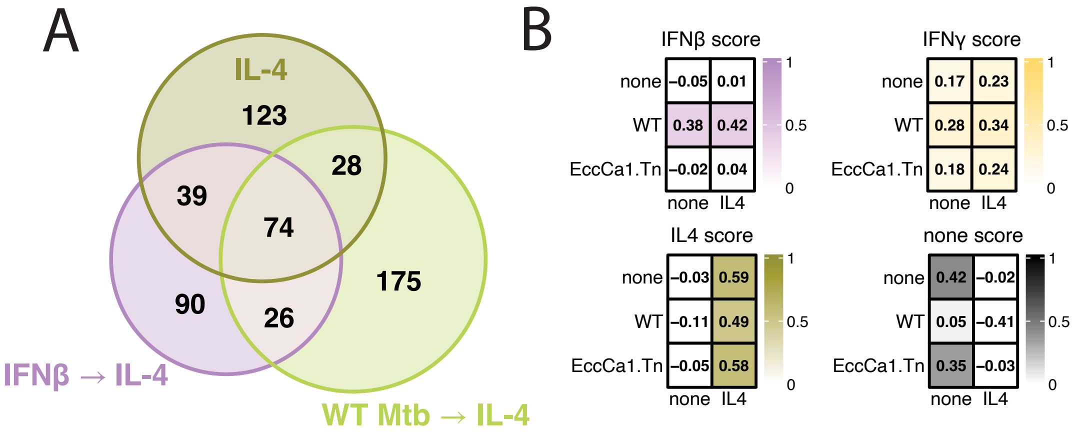

A
IL-4
123
28
39
74
90
26
175
IFNβ → IL-4
WT Mtb → IL-4
B
IFNβ score
none -0.05 0.01
WT 0.38 0.42
EccCa1.Tn -0.02 0.04
none IL4
IFNγ score
none 0.17 0.23
WT 0.28 0.34
EccCa1.Tn 0.18 0.24
none IL4
IL4 score
none -0.03 0.59
WT -0.11 0.49
EccCa1.Tn -0.05 0.58
none IL4
none score
none 0.42 -0.02
WT 0.05 -0.41
EccCa1.Tn 0.35 -0.03
none IL4
1
0.5
0
